# Tet-dependent 5-hydroxymethyl-Cytosine modification of mRNA regulates axon guidance genes in *Drosophila*

**DOI:** 10.1101/2023.01.03.522592

**Authors:** Badri Nath Singh, Hiep Tran, Joseph Kramer, Elmira Kirichenko, Neha Changela, Fei Wang, Yaping Feng, Dibyendu Kumar, Min Tu, Jie Lan, Martin Bizet, François Fuks, Ruth Steward

**Affiliations:** Waksman Institute, Rutgers University, Piscataway, NJ 08854; Department of Molecular Biology and Biochemistry, Cancer Institute of New Jersey, Rutgers University; Department of Pathology and Laboratory Medicine, Rutgers Biomedical and Health Sciences, Rutgers University, New Brunswick; Laboratory of Cancer Epigenetics, Faculty of Medicine, ULB Cancer Research Center (U-CRC), Université Libre de Bruxelles (ULB), Brussels, Belgium; Institute for Genetics, Justus-Liebig University Giessen, 35392 Giessen, Germany

## Abstract

Modifications of mRNA, especially methylation of adenosine, have recently drawn much attention. The much rarer modification, 5-hydroxymethylation of cytosine (5hmC), is not well understood and is the subject of this study. Vertebrate Tet proteins are 5-methylcytosine (5mC) hydroxylases and catalyze the transition of 5mC to 5hmC in DNA. These enzymes have recently been shown to have the same function in messenger RNAs in both vertebrates and in Drosophila. The *Tet* gene is essential in Drosophila as Tet knock-out animals do not reach adulthood. We describe the identification of Tet-target genes in the embryo and larval brain by mapping one, Tet DNA-binding sites throughout the genome and two, the Tet-dependent 5hmrC modifications transcriptome-wide. 5hmrC modifications are distributed along the entire transcript, while Tet DNA-binding sites are preferentially located at the promoter where they overlap with histone H3K4me3 peaks. The identified mRNAs are preferentially involved in neuron and axon development and Tet knock-out led to a reduction of 5hmrC marks on specific mRNAs. Among the Tet-target genes were the *robo2* receptor and its *slit* ligand that function in axon guidance in Drosophila and in vertebrates. *Tet* knock-out embryos show overlapping phenotypes with *robo2* and both Robo2 and Slit protein levels were markedly reduced in Tet KO larval brains. Our results establish a role for Tet-dependent 5hmrC in facilitating the translation of modified mRNAs primarily in cells of the nervous system.

## Introduction

The regulatory function of epigenetic mechanisms such as modifications of specific DNA bases or amino acids in histone tails have been investigated for many years. These processes are overlayed upon the genetic code and have profound effects on transcription and overall gene expression. The importance of similar modifications of RNA bases has become apparent and its pervasiveness has engendered the nascent field of epitranscriptomics [1]. Approximately 150 modifications of all four nucleosides have been detected in total RNA samples [2]. These modifications are mostly associated with the more abundant ribosomal and transfer RNAs but are also present in a subset of messenger RNA. The mRNA modifications provide a critical layer of regulation of the transcriptome in both Drosophila and vertebrates, and influence gene expression through the control of mRNA biogenesis [3]. Cytosine bases convey epigenetic information in both DNA and mRNA. 5-methylcytosine (5mrC) is present in cytoplasmic and mitochondrial ribosomal RNA, t-RNA, non-coding RNA, and mRNA [4]. In contrast, 5hmrC is most abundant in mRNA and is detected at a significantly lower frequency than 5mrC.

In Drosophila DNA, 5mC is present at low levels and so far, no function has been documented for it [5]. However, both 5mrC and 5hmrC are present in *Drosophila* RNA. The 5hmrC modification appears to be specific to mRNA and is controlled, at least in part, by the Drosophila Tet (Ten-Eleven-Translocation) protein [6]. Tet proteins were first identified as DNA-modifying enzymes that function as 5-methylcytosine (5mC) hydroxylases, catalyzing the transition of 5mC to 5hmC in vertebrate DNA [7].

The three vertebrate *TET* genes (*TET1, 2* and *3)* function as epigenetic regulators of gene expression. The transition of 5mC to 5hmC leads to the elimination of the methyl mark on DNA and activates the transcription of target genes [7]. Mammalian TET proteins, TET3 in particular, catalyze the same reaction on RNA, converting 5mrC to 5hmrC in tissue culture and mouse embryonic stem cells (ESCs) [8]. Vertebrate TET1 and TET3 isoforms have an N-terminal DNA binding domain (CxxC) and a C-terminal metal-binding catalytic domain (HxD), while TET2 lacks the N-terminal domain [9]. *Drosophila* has only one *Tet* gene, that encodes the two major protein forms from two distinct promoters [10]. The larger protein (Tet-L) includes the DNA binding and catalytic domains, while the smaller form (Tet-S) has only the catalytic domain. Both the DNA binding and catalytic domains of *Drosophila* Tet are highly conserved [11].

Complete loss-of function of *Tet* (*Tet^null^*) leads to lethality in the late pupal stage, with partial loss-of-function alleles surviving as adults for varying amounts of time[10, 12]. All mutant animals show abnormal locomotion and knock-down of Tet in neurons that control the circadian rhythm results in perturbation of that rhythm, indicating that Tet is likely essential in diverse neuronal cells. The neuronal phenotypes agree well with the expression of the Tet gene. The gene is first expressed in three-hour old embryos and persists throughout embryogenesis and larval development predominantly in the nervous system [10, 13].

Here we address the function of Tet and 5hmrC modification of mRNA which appears to occur independently of the reported 5mC to 5hmC transition in vertebrate DNA and the methylation of N6-mA in Drosophila DNA [12]. Few studies concerning the requirement of Tet in mRNA modification have been published. In tissue culture RNA modification under the control of Tet2 has been shown to lead to myeloid cell expansion through 5hmrC-based regulation of mRNAs in response to pathogen challenge [14]. Additionally, in mouse Embryonic Stem Cells (ESC), Tet proteins control the 5 hydroxymethylation of key-pluripotency transcripts as well as endogenous retroviruses [15, 16].

While Tet function in RNA modification has been analyzed in immortalized cells in *Drosophila* and mouse, we report our work on identifying genes that are regulated by Tet in *Drosophila* embryos and nerve tissue. These Tet-target genes were identified through genome- and transcriptome-wide experiments, namely ChIP-seq, hmeRIP-seq, and RNA-seq. Two of these target genes, *robo2* and *slit*, are known for their requirement in axon guidance in both vertebrates and Drosophila and we chose them for further analyses. We found that *Tet* mutant animals show overlapping phenotypes with *robo2* in the developing nervous system and that Tet activity is required for the proper expression of these pathfinding genes since loss of Tet results in reduced protein expression.

## Results

### Tet functions as a 5-methylcytosine hydroxylase and modifies polyA^+^ RNA in S2 cells, embryos, and larval brains

Previously we have shown by dot blot analysis in S2 Drosophila cells and larval brains that the 5hmrC modification was primarily found on polyA^+^ RNA and was strongly reduced in Tet knock-down (KD) cells as well as in larval brains from complete loss-of-function animals (*Tet^null^*) [6]. We have confirmed and quantified these results using ultra-high-performance liquid chromatography tandem mass spectrometry (UHPLC-MS/MS). Measurements of 5mrC and 5hmrC abundance in S2 cells indicate that 5hmrC was strongly enriched in polyA^+^ RNA whereas 5mrC was underrepresented in that fraction as compared to total RNA (Fig. 1A and B). Thus, our results are consistent with the observation that 5mrC is associated with rRNA, tRNA and polyA^+^ RNA, while 5hmrC is primarily found in mRNA, albeit at much lower levels than 5mrC. We then examined changes of 5hmrC and 5mrC in polyA^+^ RNA isolated from normal and *Tet^null^* larval brains. We found that 5hmrC was decreased about 5-fold in the mutant brains as compared to control (Fig. 1C). Moreover, 5mrC was observed to increase almost 3-fold in the absence of Tet function. (Figure 1D). Similar results were found in wildtype (wt) and Tet KD embryos (Fig. S1A and B). These results confirm and extend our previous antibody-based analyses and indicate that Tet is responsible for much of the conversion of 5mrC to 5hmrC in *Drosophila* mRNA [6].

**Fig 1.**
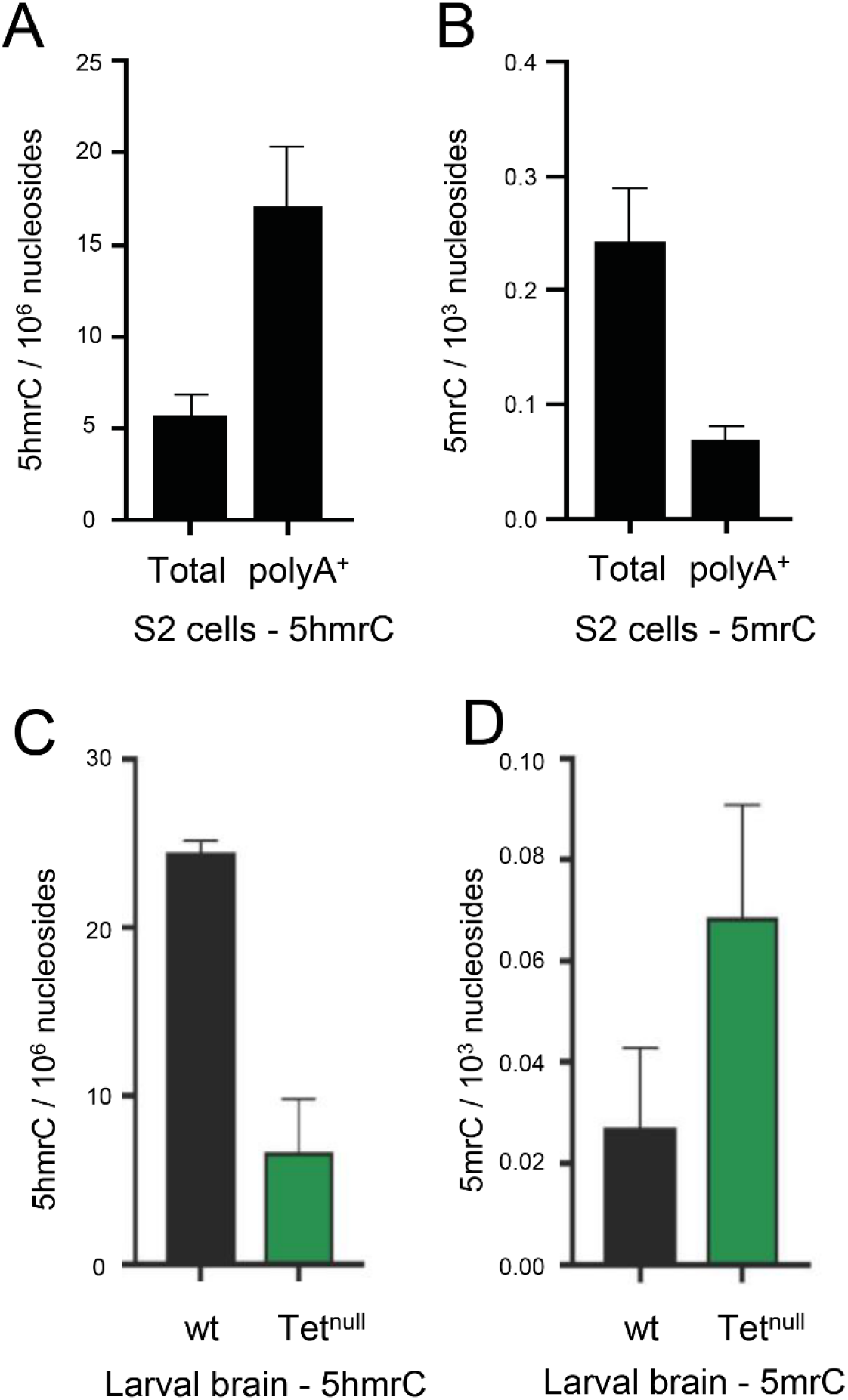
5hmrC is found in PolyA^+^ RNA and is controlled by Tet as measured by mass spectrometry. **A.** 5hmrC in total and polyA^+^ RNA isolated from S2 cells; **B.** 5mrC in total and polyA^+^ RNA isolated from S2 cells; **C.** 5hmrC in total RNA isolated from wild-type and *Tet^null^*larval brain; **D.** 5mrC in total RNA isolated from wild-type and *Tet^null^*larval brain.

### Tet binds DNA preferentially at the transcription start site of target genes

Members of the Tet protein family are known DNA and RNA binding proteins. Moreover, in vertebrates Tet proteins have been shown to bind DNA at promoter regions to regulate gene expression through active DNA demethylation [16, 17]. We sought to identify the genes that are regulated by *Drosophila* Tet. We began our experiments by determining if *Drosophila* Tet also binds DNA and mapping the binding sites. We performed ChIP-seq experiments and mapped Tet-binding peaks genome wide using chromatin isolated from the fly line that expresses the Tet-GFP fusion protein under the endogenous promoter[10]. We used two samples from different stages of development: 3^rd^ instar larval brain and imaginal discs (larval brain fraction, LBF) and 0-12h embryos. Samples were normalized to input chromatin. As negative control we used chromatin from LBF and 0-12 h embryos lacking GFP. Underlining the specificity of the anti-GFP antibody, the ChIP material from the negative control was too low to allow library preparation and sequencing (see methods).

Bioinformatic analysis of the LBF ChIP-seq results identified 3413 Tet binding peaks distributed on 2240 genes. An example of Tet binding peak profile is shown in Fig. 2A. Tet preferentially occupies promoter regions (Fig. 2B) and shows the strongest binding to promoter regions. (Fig. 2C). Analysis of the Tet bound sites identified a highly conserved CG-rich sequence via MEME-ChIP Motif Analysis (Fig. 2D and Fig. S2C). This motif is similar to that identified from similar studies of Tet1-bound loci in ESCs. [17]

**Fig 2.**
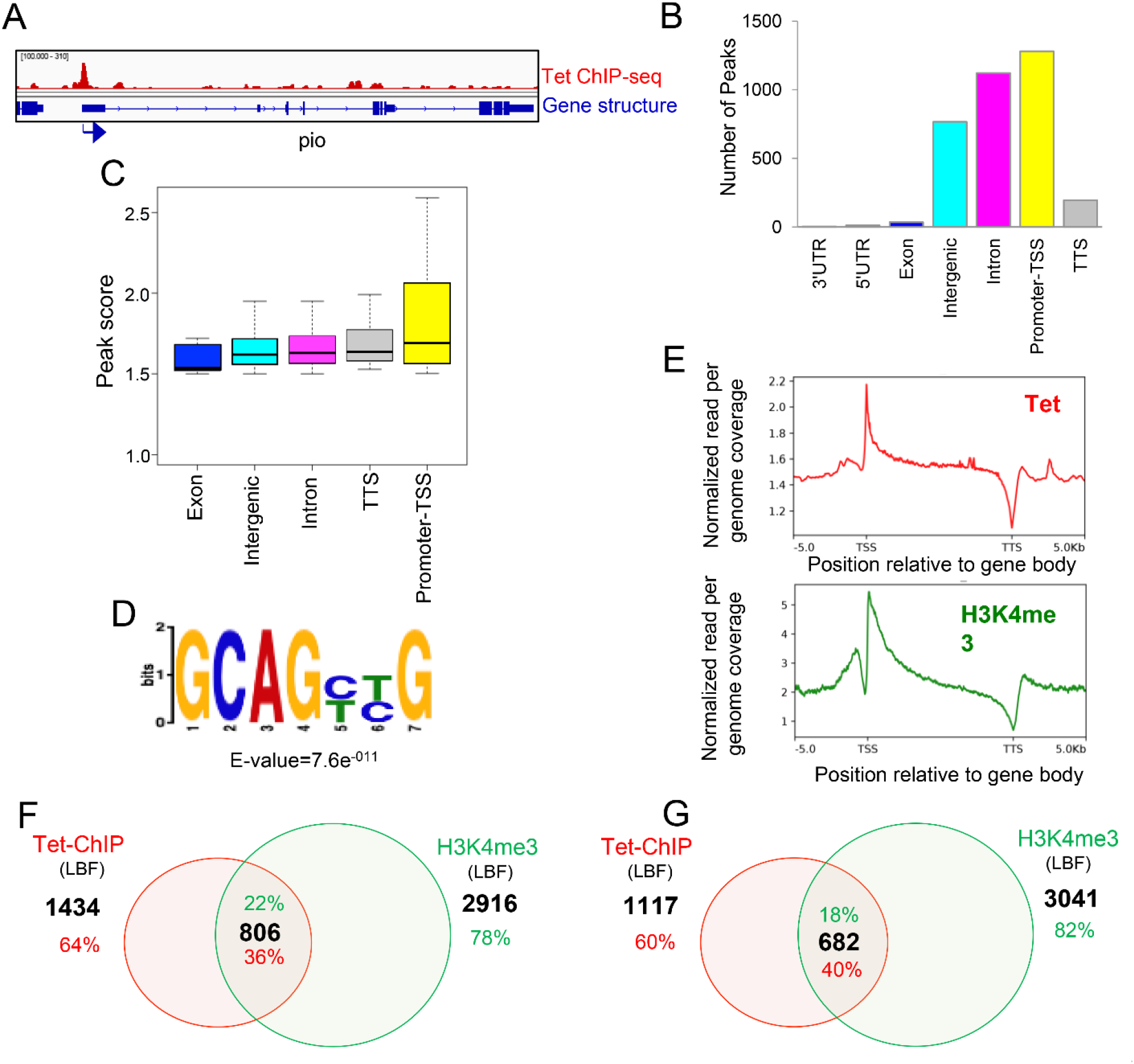
Genome-wide Tet protein binding sites in Drosophila larval brain fractions, Tet-ChIP-seq analysis: **A.** Representative gene showing Tet binding peak at the promoter. Arrow indicates promoter orientation; **B.** Genome wide distribution of Tet occupancy in larva brain fraction. The genomic regions (3’UTR, 5’UTR, exons, intergenic, introns, promoter-TSS transcription start sites, and TTS, transcription termination sites) were defined based on RefSeq gene (dm6) annotations; **C.** Strength of Tet enrichment on fly genome counted as peak score across the gene body plotted from 3413 peaks; **D.** Genome wide distribution of Tet binding sites displayed as enriched sequence motif among 3413 peaks identified by de novo motif discovery in this study; **E.** Binding profile of LBF Tet (red) and H3K4me3 (green) within the gene body ± 5kb; **F.** 36% of Tet occupied genes on various genomic regions overlapped with the H3K4me3 mark; **G.** Promoter-associated Tet binding peaks on 40% of genes overlap with H3K4me3 marks.

The composite model of Tet-binding across the coding region illustrates that Tet occupancy is highest near the promoter and gradually decreases until it undergoes a notable drop at the transcription termination sites (TTS). This closely mirrors the profile observed for H3K4me3, an epigenetic mark associated with actively transcribing regions frequently found at transcription start sites [18] (Fig. 2E). While 36% of all Tet peaks co-localize with this chromatin modification (H3K4me3, Fig. 2F), 40% of the Tet binding sites at the promoter co-localized with the H3K4me3 mark (Fig. 2G).

In embryo samples, we detected 5180 Tet-binding peaks associated with 2578 genes. An example of a Tet binding peak profile is shown in Fig. 3A. Tet is enriched throughout the gene body and intronic regions (Fig. 3B) however the strength of binding is, like in LBF, strongest at promoters (Fig 3C). The Tet-binding profile across the coding regions is similar to that observed in LBF (Fig. 3E). Analysis of the DNA sequences bound by Tet protein in embryos uncovered a highest ranking binding motif that shows significant similarity to the larval Tet consensus sequence (Fig. 3D and S2) and, as with the larval ChIP samples, we observe Tet occupancy to be correlated with H3K4me3 binding sites, at promoters (Fig 3E): 42% of all embryonic Tet peaks co-localized with H3K4me3 chromatin modification marks (Fig. 3F) and 51% of the promoter binding sites overlapped with H3K4me3 mark (Fig. 3G). In both embryos and LBF, Tet binds to approximately the same number of target genes and 30% of Tet’s targets are identical in both tissues (Fig 3H).

**Fig 3.**
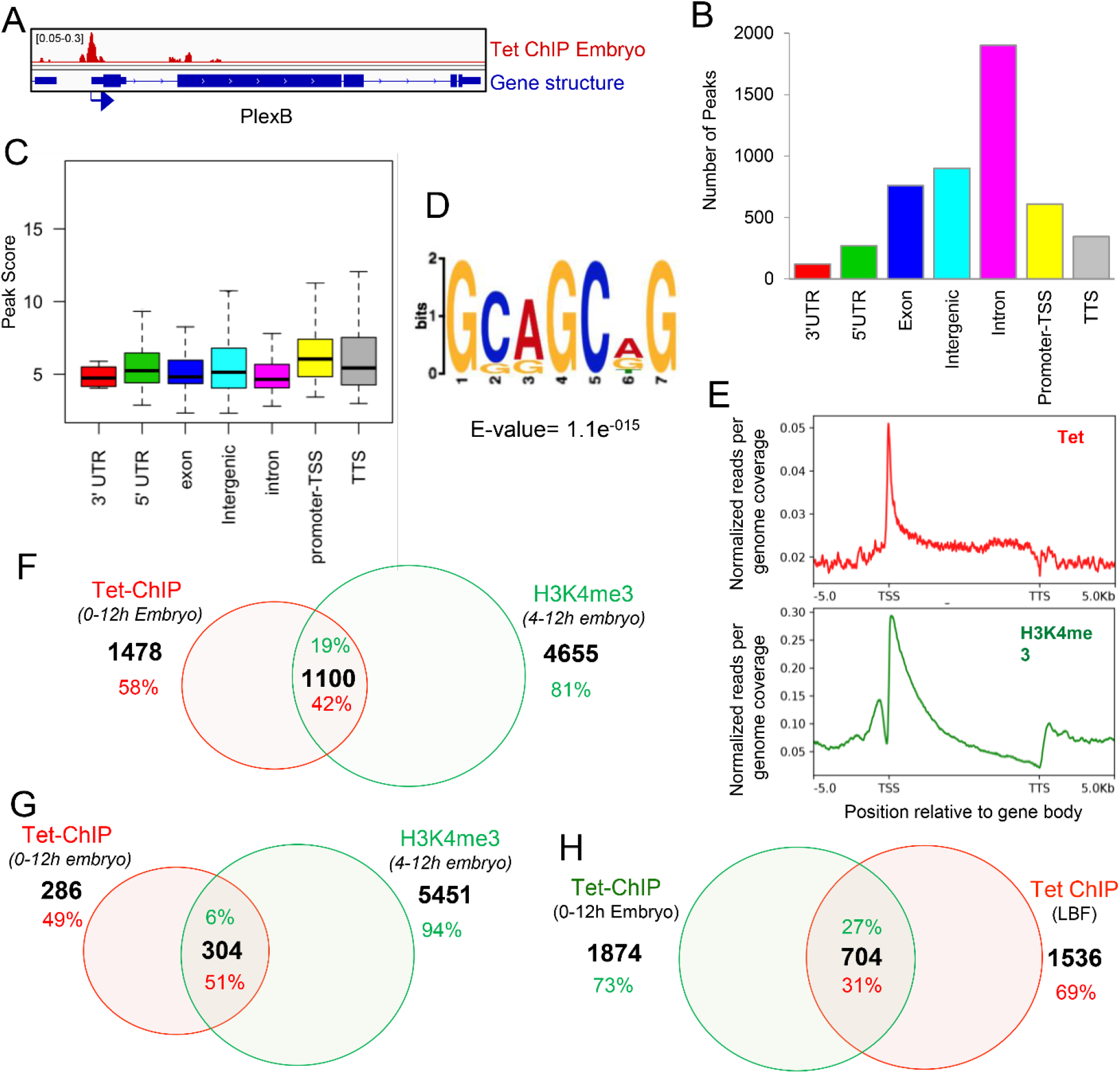
Genome-wide Tet protein binding sites in Drosophila 0-12 hour embryos, Tet-ChIP-seq analysis: **A**. Representative gene showing Tet binding peak at the promoter. Arrow indicates promoter orientation; **B.** Genome wide distribution of embryo Tet ChIP-seq peaks in different genomic regions; **C.** Strength of Tet enrichment on different genomic regions counted as peak score plotted from 5180 peaks; **D.** Enriched sequence motif among 5180 embryo Tet ChIP-seq peaks identified by de novo motif discovery in this study; **E.** Binding profile of embryo Tet (red) and H3K4me3 (green) within the gene body ± 5kb; **F.** 42% of Tet bound genes in embryo have H3K9me3 modification; **G.** 51% of genes that show binding of Tet to the promoter that overlap with H3K4me3; **H.** 27% of Tet bound genes in embryo also have Tet binding peaks in larva brain fraction.

Our ChIP-seq results indicate that Tet binding sites are distributed throughout the physical map of the genome (Figs S2A and S2B). To confirm these results and show that the Tet-DNA binding domain is sufficient to target Tet to DNA, we constructed transgenic flies carrying a Myc-tagged DNA-binding domain of Tet (CxxC) under the control of the heat shock promoter (hsp70-GAL4::UAS-TetCxxCRFPmyc). We expressed the Tet DNA-binding domain by exposing larvae to heat shock and stained salivary glands with anti-Myc and anti-H3K4me3 antibody. As indicated by Chip-seq, Tet showed many bands distributed on all arms of the chromosomes, but virtually no staining of the chromocenter which contains very few genes. H3K4me3 is also present in a distinct binding pattern on all chromosomes, but in contrast to Tet is abundant in the chromocenter and the nucleolus. These staining results agree with our observation that Tet binds to genes on all chromosomes of *Drosophila* (Fig S2A).

Our Chip-seq experiments were done in embryos and LBF, two tissues at diverse stages of fly development, but in which Tet protein is highly expressed. In both tissues we identified about 2500 genes that showed significant Tet-binding genome-wide. Tet binding characteristics were similar in both tissues in that the most significant Tet-binding peaks, showing strongest binding, were preferentially located at promoters. About 30% of the Tet DNA-binding sites are identical in embryos and LBF. Thus, it appears that only some of the Tet targets are fixed while others show stage-specific variations throughout development.

### Identification of Tet-target mRNAs by hMeRIP-seq in fly tissues

We next determined how many of the genes with Tet-binding peaks also showed 5hmrC modifications of their RNA. To do this we mapped Tet-dependent 5hmrC modifications on RNAs transcriptome-wide in the same tissues we used for our Chip-seq analysis. We first performed hMeRIP-seq on total RNA using basically the same approach we used previously in S2 cells [6]. RNAs isolated from wt 0-12 h embryos and from wt and *Tet^null^* Larval Brain Fractions (LBF) were treated with anti-5hmC antibody or immunoglobulin as negative control, and followed by Next Generation Sequencing (NGS, see methods).

In the embryo, we identified 1815 peaks on 1402 mRNAs. A representative 5hmrC peak profile is shown in Fig. 4A. The 5hmrC modification is preferentially associated with coding sequences and a comparison to the expected distribution of peaks shows that the distribution of the modification is not random (Fig. 4B). Moreover, as the presence of the 5hmrC modification is not proportional to the abundance of the mRNA the modification appears to function broadly within the transcriptome and is not a regulatory modality restricted either to rare or hyperabundant transcripts (Fig. 4C). The 5hmrC-associated sequences identified from these experiments revealed a specific UC-rich motif present within these mRNAs that closely resembles the motif observed in S2 cells and mammalian ESCs (Fig. 4E and Fig. S3) [6, 16].

**Fig 4.**
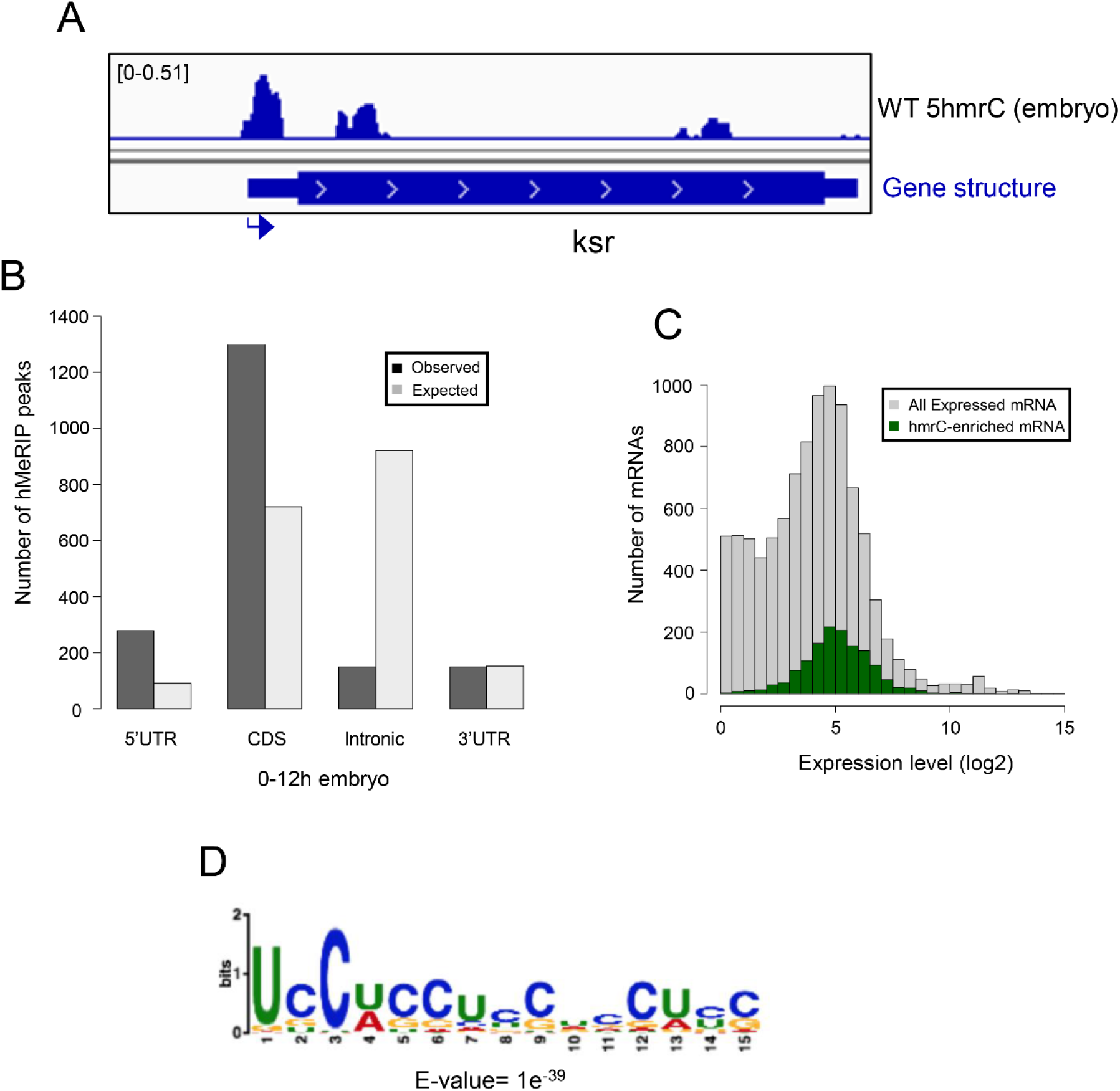
Transcriptome-wide distribution of 5hmrC in Drosophila 0-12 h embryo mRNA, hMeRIP-seq: **A**. Example of gene showing 5hmrC peak distribution. Arrow indicates promoter orientation; **B.** Distribution of 5hmrC peaks on embryonic transcripts and comparison of actual and predicted peaks according to the type of structural element within the transcript; **C.** Distribution of all expressed (gray) or 5hmrC enriched (green) transcripts, showing the number of mRNAs as a function of their expression levels in wt embryo; **D.** Sequence motif identified in within 1815 5hmrC peaks.

In mRNA from the wild type LBF, we detected 3711 peaks on 1775 transcripts. A representative profile of 5hmrC enriched peaks in wt and *Tet^null^*is shown in Fig. 5A. In wt the peaks were distributed across the gene body (Fig. 5B) and 5hmrC marks were found to decorate mRNAs independent of their abundance (Fig. 5D). Analysis of the peak sequences indicated the modifications were primarily associated with a UC-rich motif highly related to that identified in embryonic samples (Fig. 5F). In mRNA from *Tet^null^* larvae we identified 5,374 peaks in 1710 mRNA. Comparison of mRNAs identified in both the wt and *Tet^null^* samples indicate that the distribution of 5hmrC peaks is similar both in the presence and absence of Tet function. However, In the *Tet^null^* samples, 45% of the transcripts identified had at least one peak that showed a reduction in the 5hmrC modification relative to wild-type (Fig. 5C) and the reduction was most pronounced on intronic and coding region peaks (45% and 46%) compared to the peaks found in the UTRs (5’, 19%, and 3’, 16%). Thus, within a given mRNA transcript some peaks were affected in *Tet^null^* LBF, while others remained unchanged. These results suggest the preference of Tet to modify specific regions of transcripts.

**Fig 5.**
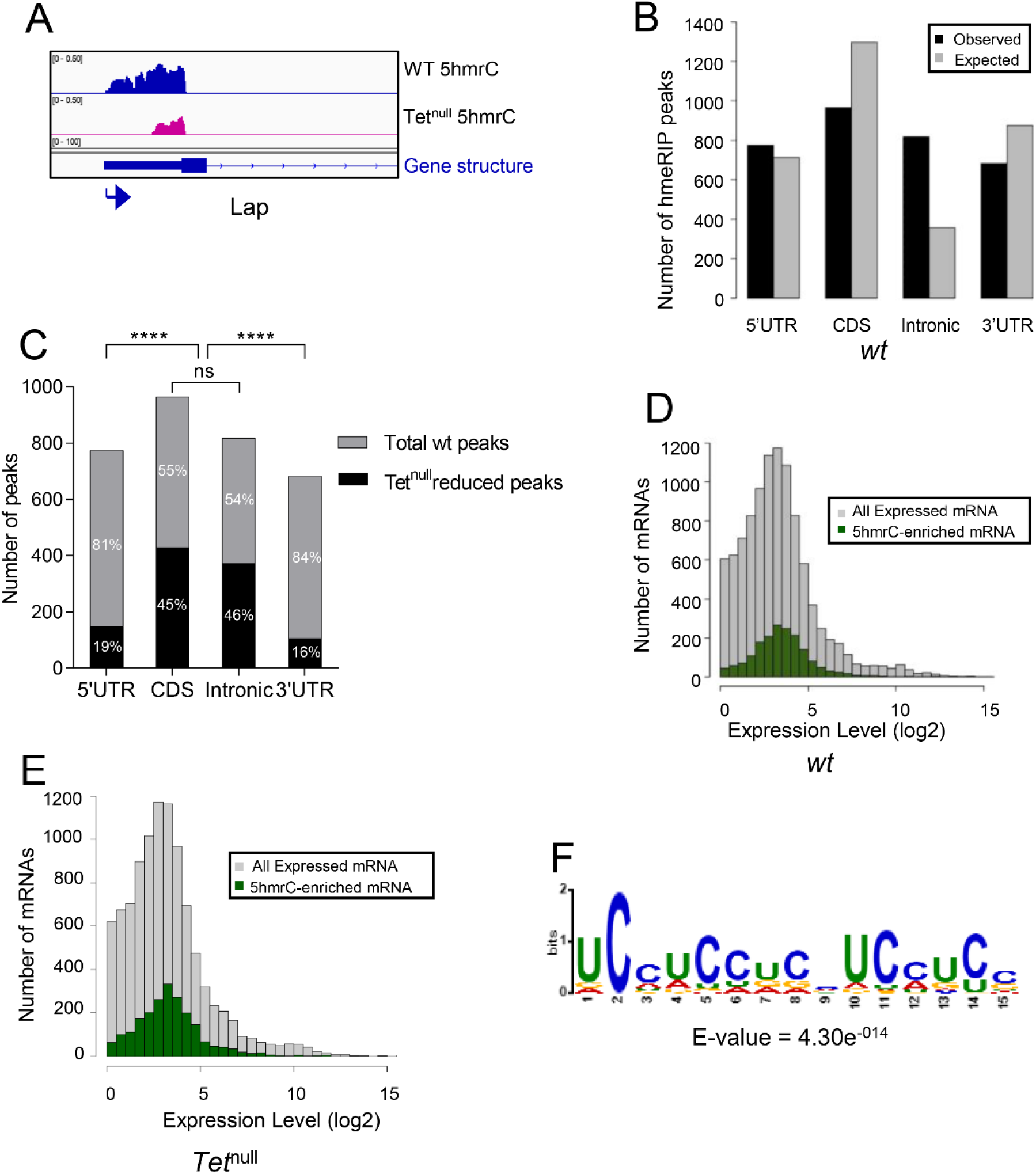
Transcriptome-wide distribution of 5hmrC in LBF mRNA, hMeRIP-seq: **A**. Example of gene showing 5hmrC peak distribution. Arrow indicates promoter orientation; **B.** Distribution of 5hmrC peaks on wt LBF transcripts and comparison of actual and predicted peaks according to the type of structural element within the transcript; **C.** Distribution of 5hmrC peaks reduced in *Tet^null^* compared to the peaks found in the wt LBF; note that peaks in the protein coding sequences and introns are significantly more reduced in *Tet^null^* than are the peaks in the 5’ and 3’ UTR; **D.** Distribution of all expressed (gray) or 5hmrC enriched (green) transcripts, showing the number of mRNAs as a function of their expression levels in wt LBF; **E.** Distribution of all expressed (gray) or hmrC enriched (green) transcripts, showing the number of mRNAs as a function of their expression levels in *Tet^null^* LBF; **F.** Sequence motif identified within 3711 5hmrC peaks.

In addition, 37% of the modified mRNA in embryos were also identified in the LBF, while 30% of the larval modified mRNAs were also present in the embryonic fraction (Fig. S4C). Taken together these results suggest that Tet targets a distinct cohort of mRNAs in embryos and larval brains and controls specific 5hmrC modifications along transcripts.

### RNA levels in wild type and *Tet^null^* larval brains

Our results indicate that Tet binds to the promoter of a subset of possibly actively transcribed genes and controls the 5hmrC modification of their mRNAs. The modification may have an effect on the stability, processing, and/or translation of the transcripts. To determine if there is a link between 5hmrC modification and mature mRNA levels, we sequenced (NGS) RNA isolated from wildtype and *Tet^null^* LBF. We found that out of 9000 total transcripts the levels of 445 were significantly increased and 115 were decreased in *Tet^null^* LBF (Fig. 6A). When we compared these mRNAs with the 5hmrC-modified mRNAs present in LBF, we found that 1716 or ∼20% of the total transcripts were modified. However, of these modified mRNAs only 15, or 3 % were upregulated in *Tet^null^*, and 13 or 11 % were decreased (Fig. 6A, B). This result indicates that the levels of the vast majority of 5hmrC modified mRNAs do not change levels in *Tet^null^*LBF. Thus, the 5hmrC modification of the mRNAs does not appear to generally control the steady state level of transcripts.

**Fig 6.**
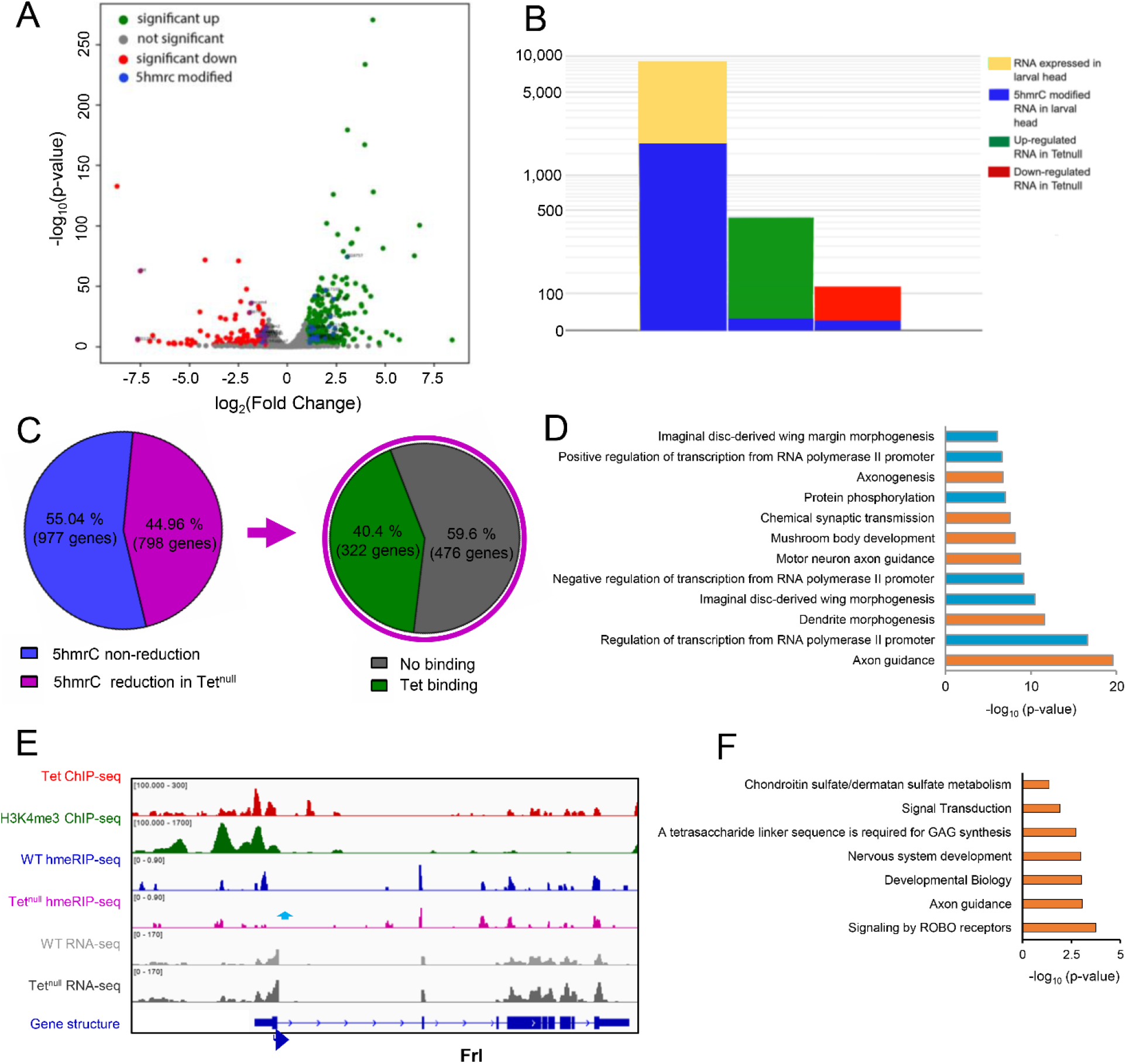
The 5hmrC modified mRNAs. **A.** Volcano plot of mRNAs that are increased (green) or decreased (red) relative to wildtype levels in *Tet^null^* LBF preparations; **B.** Proportion of modified mRNAs in all 9000 wild type transcripts, and in the decreased and increased portions of mRNAs from *Tet^null^* LBF; note the low level of modified transcripts in these two groups of mRNAs; **C.** The percent of transcripts that show a reduction of 5hmrC modification in Tet^null^ compared to wt and the percent of transcript that showed both 5hmrC reduction and Tet binding to the corresponding gene; **D.** GO term analysis of transcripts that show reduction in 5hmrC modification; **E.** IGV tracks of a representative gene showing the distribution of indicated peaks along the gene body; **F.** Pathway analysis of neuronal genes shown in **D**. ChIP-seq, hMeRIP-seq and RNA-seq data are shown in reads per million with the y-axis. Genomic regions with statistically significant enrichment were measured by -log10 (peak P values); P<10^-8^) are indicated. The effects of Tet depletion on 5hmC levels are also represented. The Y axis scale is indicated above each track. Blue arrows show reduction in 5hmrC peaks.

### Cellular function of genes controlled by Tet

Tet protein is detected in embryos from blastoderm stage onwards and is most strongly expressed in neuronal tissues and also in cardiac and muscle precursor cells. In third instar larvae, the gene is strongly expressed in the brain and neuronal cells in imaginal discs [10]. It was therefore important to assess if our molecular analyses would agree with this expression pattern and if target genes are associated with neuronal functions. We performed Gene Ontology (GO) analyses of the genes identified via ChIP-seq as well as of the genes encoding the 5hmrC-modified mRNAs that were identified in our hMeRIP-seq analyses in the embryo and the LBF (Fig. S5 A-D). The genes identified in both embryonic and larval samples through both ChIP-seq and hMeRIP-seq all show enrichment for genes involved in axon guidance. When we looked at the GO terms of transcripts that showed a reduction of the 5hmrC modification in *Tet^null^* samples, axon guidance genes were highly represented, in fact, GO terms of transcripts showing reduction of the modification in *Tet^null^* samples identified mostly genes associated with neuronal functions (see highlighted genes in Fig. 6C). Pathway analysis showed that ROBO receptor signaling is the most enriched pathway in the group of neuronal genes (Fig. 6F).

It is striking that in our two very different experimental approaches, ChIP-seq and hMeRIP-seq we identified genes with overlapping functions (Fig. S5 A-D). The importance of our results is also underlined by the observation that of the transcripts that reduction of 5hmrC levels in *Tet^null^* samples, 40% were derived from genes that also have at least one Tet DNA-binding site (Fig. 6C). In LBF samples, 43% of all the transcripts that show 5hmrC modification are derived from genes that have been shown to bind Tet (Fig. S4A). In embryo samples, 29% of all the transcripts that showed 5hmrC modification are derived from genes that bind Tet (Fig. S4B). Further, 29% of modified transcripts in embryos and 37% of modified transcripts in LBF show 5hmrC marks at both developmental stages (Fig. S4C). An example of the experimental IGV tracks of all our results for a gene in the larval CNS and the embryo are shown in Fig. 6D and Fig. S6A, respectively.

These analyses show that Tet-dependent 5hmrC is often found on mRNAs derived from genes that show Tet binding. Notably, close to 50% of transcripts that show a reduction in the 5hmrC mark in *Tet^null^* tissues are derived from Tet-target gene identified by ChIP-seq. However, the levels of these mRNAs are generally unaffected by the loss of Tet suggesting that the 5hmrC modification does not affect steady state level of mRNAs but other aspects of mRNA function such as translation or localization.

### Tet target genes

We used the results above to identify Tet-target genes and sought to determine whether the phenotypic effects of the loss of Tet’s activity were derived from its inability to regulate target mRNAs [6, 10]. We looked for genes that are 1. active in the nervous system where Tet is enriched and 2. showed Tet protein binding to DNA, and 3. whose mRNA showed a reduction in 5hmrC in *Tet^null^*animals. Axon guidance genes as a group frequently showed Tet-DNA-binding and 5hmrC mRNA modification by Tet (Fig. 6D) and Robo receptor signaling is the most enriched pathway. Among the genes that fulfilled the three criteria were two well-studied genes that function in Robo receptor signaling, *robo2* and *slit* (Fig. S7). The Slit/Robo signaling pathway is required for axonal pathfinding and the bilateral organization of the CNS in both vertebrates and invertebrates [19]. Robo proteins are transmembrane receptors on axonal growth cones for the secreted Slit ligands. Glial cells present at the midline secrete Slit and signaling between Robo and Slit is essential to inhibit midline crossing of axons through commissures via repulsion [20]. Importantly, Slit has previously been implicated as a target of Tet activity in midline glia [13]. We examined axonal pathfinding in the embryonic ventral nerve cord (VNC) and reasoned that if Tet impinges upon the levels of Robo2 and/or Slit, we should observe midline defects in *Tet^null^* animals like those seen in *robo2* or *slit* mutant embryos. Gross CNS commissural structure is maintained in *Tet^null^* embryos (Fig. 7B’, HRP), however, examination of neuronal subpopulations within the longitudinal neuropils indicates frequent pathfinding defects. A well described neuronal subpopulation, Fas2+ neurons, exhibit extensive midline crossing of growth cones in these *Tet^null^* embryos (Fig. 7B, arrows; Table S1). Additionally, the most lateral of the Fas2^+^ longitudinal tracks are often incomplete or absent (Fig. 7B, 46%-arrowheads). A second subpopulation of neurons expressing Connectin also appears to be altered in *Tet^null^* VNCs and fails to populate one of the longitudinal tracks compared to wild type (Fig. S7B; arrows). These phenotypes are strikingly similar to the axonal pathfinding defects seen in *robo2* embryos with Tet’s effects being slightly more severe (Fig. 7B and C and Table S1) [20]. We sought to determine whether the reduction of Tet-mediated 5hmrC deposition on the *robo2* or *slit* mRNAs resulted in mRNA species with reduced activity or potential for expression. Thus, we examined genetic interactions between Tet and the Slit/Robo signaling pathway in *Tet^null^* embryos lacking one copy of *robo2* or *slit*. We additionally examined Robo1, a gene that is also involved in midline repulsion but is not 5hmrC modified. Decreasing the dose of Robo2 or Robo1 in a *Tet^null^* background has little effect on Fas2+ axonal pathfinding in comparison to *Tet^null^* alone (table S1). The failure to see an effect with Robo2 may stem from the observation that the levels of midline crossing in *Tet^null^* embryos exceeds that seen for *robo2*^null^ embryos (Table S1 and [21]). However, reducing the gene dose of *Slit* by half enhances the midline crossing of Fas2+ neurons in *Tet^null^* embryos (Table S1; Figure 7B and E; 48% vs 32% *Tet^null^*), whereas heterozygous *slit* embryos show midline crossing in < 1% of segments (Fig. 7D). Moreover, *Tet^null^* mutant animals appear to be sensitized towards midline crossing in general when lacking full *slit* function. Notably, the commissures (red arrowheads, Fig. 7E’) are poorly defined, likely due to too many axons inappropriately transiting the midline.

**Fig 7.**
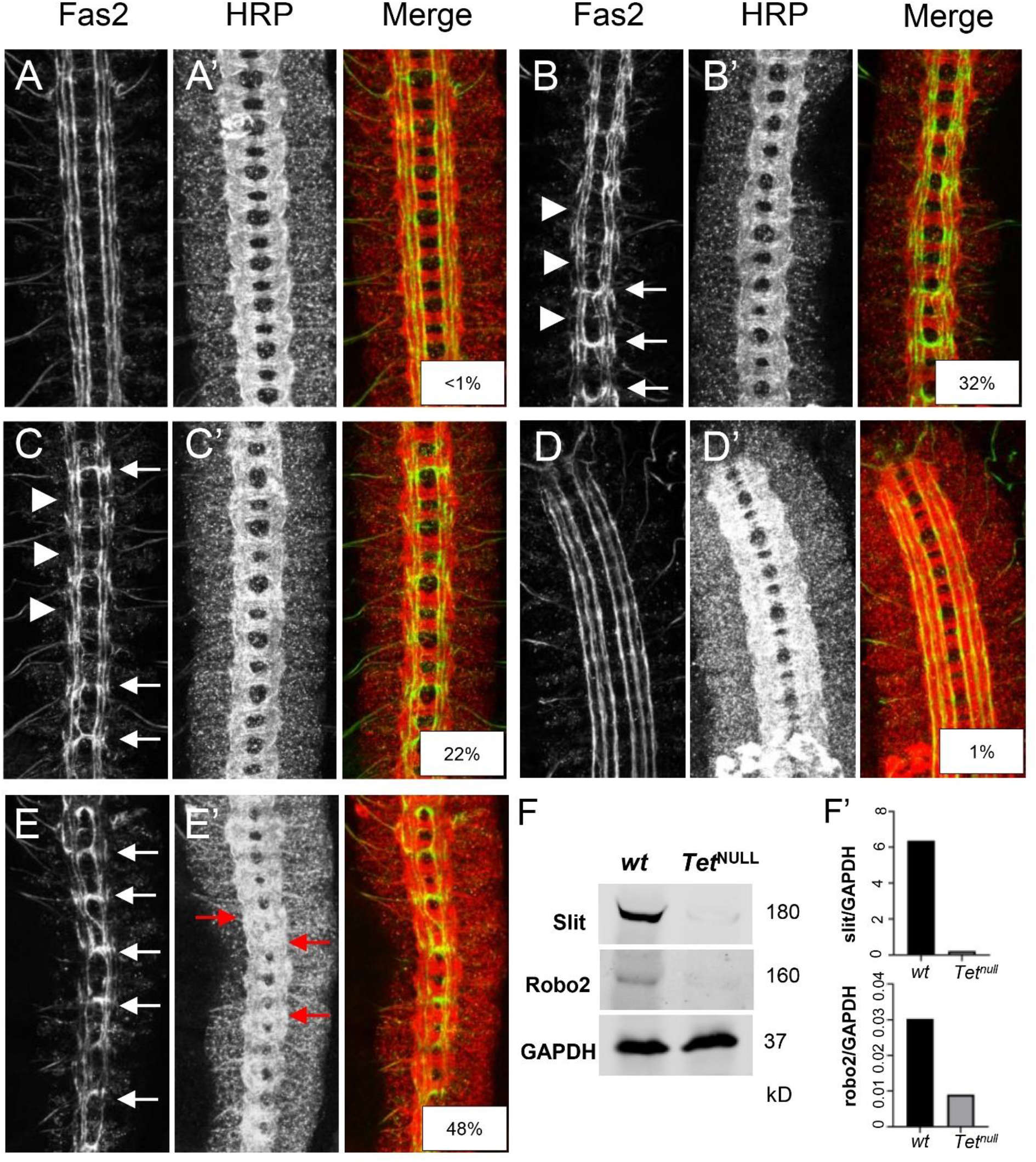
Tet regulates the expression of members of the Slit/Robo signaling pathway. Stage 16/17 embryonic ventral nerve cords immunolabelling a subpopulation of CNS neurons with Fas2 (**A-C)** and the general neuronal cell surface marker, HRP (**A’-C’)**. **A, A’**: wild-type; **B, B’**: *Tet^null^/Tet^null^*; **C, C’**: *robo2^x123^/robo2^x123^*; **D, D’**: *sli^2^*/+**; E, E’**: *sli^2^*/+; *Tet^null^/Tet^null^*. Examples of midline crossing are indicated by white arrows and malformed lateral Fas2 tracks are noted with white arrowheads. Red arrows in **E’** highlight commissural malformations present in *Tet^null^/Tet^null^* embryos with reduced slit dosage. Percentage midline crossing is displayed in the overlay panels. **F.** Western blot showing Slit and Robo2 proteins in *wt* and *Tet^null^/Tet^null^*3^rd^ Instar larval brain extracts. GAPDH is the loading control; **F’.** Normalized levels of Slit and Robo2 quantitated via optical densitometry.

**Fig 8.**
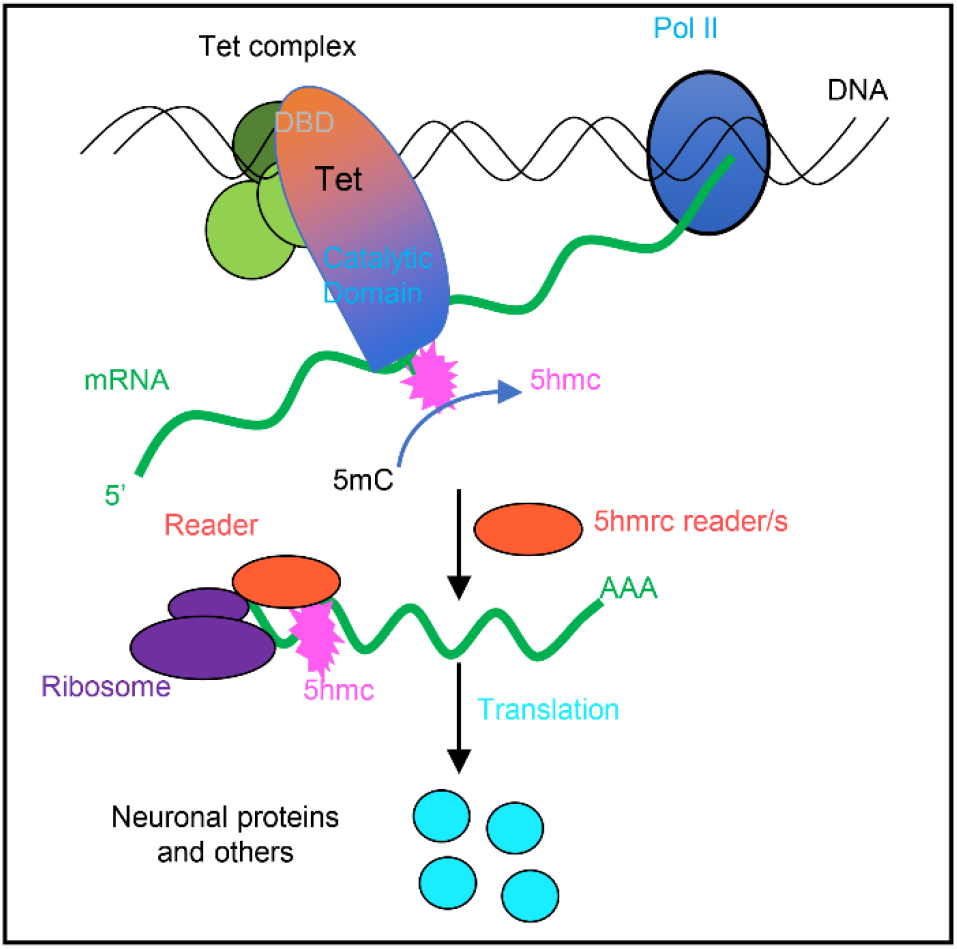
The proposed model of Tet functions in RNA modification (see text for description). Based on all our results we suggest the model shown in Figure 8, we propose that Tet binds, possibly as a complex to DNA binding sites mediated by its DNA-binding domain. The Tet binding sites are preferentially located at promoter regions of genes that also show H3K4me3, generally accepted as a mark of active transcription. We further postulate that Tet binds nascent mRNA through its RNA binding domain or possibly in cooperation with associated proteins (RNA-binding proteins, and with a so far unidentified RNA methyltransferase) to set the 5hmC mark. The 5hmrC marked mRNAs are then exported from the nucleus and recognized by a reader protein that will control the efficient loading of the modified mRNAs onto polysomes, where the mRNAs are proficiently translated.

Given that *robo2* or *slit* encode mRNAs that carry 5hmrC mark and exhibit a reduction in a *Tet^null^* background while maintaining normal steady state mRNA levels, we expected Tet to potentially control their protein levels (Fig. S7). Indeed, both proteins were clearly reduced in brain extracts from *Tet^null^* larvae relative to wt (Fig. 7F and 7F’). These results support the idea that one function of Tet-dependent 5hmrC modification is to control high levels of translation of specific target mRNAs and that in the context of embryonic axonal pathfinding Tet provides an additional, novel layer of regulation of the medically important Slit/Robo pathway.

While several aspects of this model need to be investigated our results provide a consistent framework of how Tet and Tet-dependent RNA modifications may function in controlling gene expression. Recently, mutations in human Tet3 have been shown to cause neurodevelopmental delays. It will be interesting to investigated if 5hmrC RNA modification is deficient in the affected patients [22].

## Discussion

In our previous study we investigated if Tet proteins, that are well known as 5-methylcytosine (5mC) hydroxylases catalyzing the change from 5mC to 5hmC in DNA, can have a similar function in RNA [6]. For these molecular studies we mainly used *Drosophila* S2 cells as source material. In the present study we used animal sources, embryos, and larval brain tissues, to investigate the function of Tet in modifying mRNA *in vivo*. We also wanted to delineate the molecular and cellular processes for which the modification is required, and to identify in vivo targets of the Tet protein.

Our results confirm the presence of the 5hmrC modification in mRNA by mass spectrometry in embryos, larval brain tissue and S2cells. We further show that Tet protein binds to DNA at distinct sites, functions in modifying mRNAs, and that this modification modulates translational output of the mRNAs. We used our molecular results to identify Tet target genes. We selected genes that, 1. contain promoter proximal Tet-binding site(s) that overlap with H3K4me3 modifications, 2. whose mRNA showed 5hmrC modifications that were reduced in *Tet^null^* neuronal tissues, and 3. whose mRNA levels displayed negligible changes in *Tet^null^* neuronal tissues.

We found that these target genes were most often associated with axonal growth and pathfinding. Two such genes, *robo2* and *slit*, were selected because they fulfill the conditions outlined above and are members of a conserved set of cell-signaling molecules responsible for controlling the activity of axonal growth cones of the developing CNS in vertebrates and invertebrates [23]. Phenotypic analysis of the developing CNS in Tet-deficient animals indicates a specific requirement for Tet in the proper patterning of the CNS; *Tet^null^* embryos showed a similar CNS phenotype to Robo2 deficient animals. Indeed, in the absence of Tet levels of Robo2 and Slit proteins are reduced in the larval brain, resulting in aberrant axonal pathfinding and other defects in nervous system patterning [10, 13].

### Tet controls the 5hmrC modification on mRNA

In mass spectrometry experiments we determined that 5hmrC is strongly enriched in polyA^+^ RNA confirming our previous dot blot results. This modification is much rarer than other well-studied mRNA modifications, such as 5mrC or 6mA (Fig. 1) [6, 24]. Because Tet is expressed in Drosophila almost exclusively in nerve cells, we determined the levels of 5mrC and 5hmrC in two tissues that show high Tet expression, wild type 0-12 h embryos and in larval brains. We found that 5mrC levels are about two orders of magnitude higher than 5hmrC levels (∼2x10^5^ 5mrC and ∼2x10^7^ 5hmrC in larval brains), and therefore detecting 5hmrC is not trivial.

The presence of 5hmrC is notably reduced (∼ 5 fold) in *Tet^null^*samples. Our results are consistent with the *Drosophila* Tet enzyme being responsible for this 5hmrC modification (Fig. 1 and S1). However, the remaining ∼20% of the wild type 5hmrC levels in mutant tissues that lack Tet, point to the presence of an additional hydroxymethyltransferase(s) that can modify 5mrC in the *Drosophila* genome. The existence of additional enzyme(s) contributing to mRNA hydroxymethylation has also been postulated in mouse ESCs [16].

Our mass spectrometry findings and the results from our hMeRIP-seq experiments on larval brain fractions (LBF) and embryos are consistent with what has been previously reported for *Drosophila* tissue culture cells and for ESCs (Fig. 1,4,5 and S1, S3) [16]. We identified ∼3000 5hmrC peaks in ∼1500 transcripts in S2 cells [6]. In ESCs the number of peaks was 1633 in 795 transcripts [16]. In our *in vivo* experiments we identified 1815 peaks in 1402 transcripts in embryos, and 3711 peaks on 1776 transcripts in LBF. Of the modified transcripts in embryos 37% were also identified as modified transcripts in the LBF. In all samples the modification peaks centered around a UC-rich consensus motif (Fig. S3). The consistency of the mapping results of the 5hmrC modifications in *Drosophila* tissue culture cells, embryos, larval brain fraction, and ESCs underlines the probable conserved function of Tet across the species.

The 5hmrC peaks on mRNAs derived from LBF are distributed all along the transcripts, the UTRs, the coding region, and introns. However, in *Tet^null^*LBF peaks in the CDS and introns are significantly more strongly reduced than peaks in the UTRs (Fig. 5C). This observation suggests that Tet may target coding sequences and introns specifically. We do not yet understand if modifications in different parts of the transcripts have diverse functions and if they may be controlled by additional enzyme(s).

### Drosophila Tet’s DNA binding activity

We found that in both embryos and in LBFs, Tet recognizes a DNA motif similar to the motif bound by Tet1 in vertebrate ESCs (Fig. S2C) [17, 25]. A majority of these peaks are associated with coding regions and are frequently found at the promoter. Almost 50% of the peaks overlap with the H3K4me3 mark, an indication that the genes are actively transcribed. The distribution of Tet-binding peaks and the overlap with the H3K4me3 mark agree well with the localization of the Tet-DNA-binding domain on salivary gland chromosomes confirming that the binding sites are found almost exclusively in euchromatin and are distributed on all 4 chromosomes (Fig. S2A).

We propose that the selection of target RNAs modified by Tet is at least in part facilitated by Tet’s DNA-binding of specific genes. The concurrence of Tet-DNA binding peaks on genes that also showed Tet-dependent 5hmrC modifications of their mRNA is consistent with this idea. The majority of the genes that show Tet binding and modified mRNAs are divergent in both tissues indicating that in addition to a conserved function of Tet in different neuronal cells, Tet also has a tissue-specific or possibly even cell-type-specific function.

### Identifying Tet target mRNAs

Tet is highly expressed in nervous tissues and the loss of Tet function leads to abnormal neuronal functions such as defects in larval locomotion or abnormalities in the circadian rhythm. [10] Our immunoprecipitation of 5hmrC-modified RNAs identified 1775 genes in larval brain fractions. 45 % (798) showed a significant decrease in the overall 5hmrC peaks in a *Tet^null^*background. Of the genes with reduced 5hmrC marks, 44% showed Tet-DNA binding. Notably, the mRNAs in which the reduction of the 5hmrC mark was seen were mostly associated with genes that function in different aspects of nerve cell development. First among them are axon outgrowth genes that were also identified in the GO-term analysis as abundant gene categories associated with Tet binding sites and mRNAs carrying the 5hmrC mark (Fig. 6D, S5).

Our initial examination of the developing embryonic ventral nerve cord (VNC) in *Tet* mutants identified subtle defects in CNS patterning. We then examined subsets of VNC neurons using antibodies to Fas2 and Connectin (Fig 7B, B’ and S7B, B’) guided by our molecular results. Overall commissural structure is maintained in *Tet^null^* embryos, however neurons expressing Fas2 show a failure of the midline to repel axon crossing effectively. And so, we looked among the Tet mRNA targets with known functions in axon guidance and found that both *slit* and *robo2* mRNAs were represented. Both genes have Tet-binding sites near the TSS, their mRNA is modified, and the modification is reduced in *Tet^null^* LBF, while their mRNA levels are not significantly changed (Fig. S7). Comparison of the CNS in *Tet^null^* and *robo2^null^* embryos identified a set of overlapping phenotypes with high frequency midline crossing defects of Fas2+ neurons, as well as discontinuities in the most lateral, longitudinal Fas2 and Connectin axonal tracts (for description of embryonic nerve cord see [20]). Notably, these tracts correspond to neurons which express the Robo2 protein [26, 27].

The overlapping phenotypes of Tet, *robo2 and slit,* together with the molecular data that identified Robo2 and Slit as Tet targets, prompted us to investigate if Robo2 and Slit protein expression was affected by the loss of Tet. Indeed, in Western blots from *Tet^null^* larval brain extracts both Robo2 and Slit protein levels were strongly reduced (Fig. 7F, F’), indicating that Tet’s profound consequences on VNC patterning occurs, at least in part through the control of expression of the Robo2 and Slit proteins. As Robo2 and slit mRNA levels are not changed in *Tet^null^* LBF (Fig. S9), we suggest that the Tet-dependent 5hmrC modification positively controls the level of translation of the two mRNAs. While we have not investigated the protein levels of additional Tet-targets, we expect that Tet controls protein levels through the 5hmC modification of many target mRNAs. Which step in RNA processing leading to mRNA translation is affected in *Tet^null^* animals will have to be elaborated. Based on our previous results, that showed that 5hmrC modified RNAs are found on polysomes, at least one possibility is that the 5hmrC modification facilitates the loading of the mRNAs on ribosomes [6].

Our work supports a function of Tet in controlling the 5hmrC modification of specific neuronal mRNAs, essential for maintaining translation levels necessary for normal neuronal function, thus adding an additional level of control of gene expression. However, we cannot exclude that Tet has additional functions in controlling gene expression in Drosophila.

## Materials and Methods

### Drosophila Genetics

All flies were reared at 25°C and kept on standard medium. The mutant Tet alleles are described in [6, 10]; the wild-type allele used in all experiments is *w^1118^*. The stock utilized to examine Robo2 was robo2^x123^/CyO [28]. The material used for all whole-genome analysis was either hand dissected third instar larval brains, or, because some experiments necessitated a large input, dissected anterior parts of larvae including the 3 anterior abdominal segments that contain the brain besides other tissues such as imaginal discs, salivary glands, mouth parts and epidermis. Because Tet is highly expressed in the brain and the nerve cell in discs, but not in the other tissues, we call this the Larval Brain Fraction, LBF. Brains and larvae from wt and Tet-GFP third instar larvae were dissected in cold-PBS supplemented with protease inhibitor, snap frozen on dry ice, and stored at -80°C.

### Immunohistochemistry and Imaging

The following antibodies were used for immunolabelling of late stage embryos and chromosomal preparations: mouse anti-Fas2 (Developmental Studies Hybridoma Bank, DSHB), rabbit anti-HRP (Jackson Immunoresearch), mouse anti-Connectin (DSHB), rabbit anti-dsRED (Invitrogen), rabbit and mouse anti-GFP (Invitrogen), mouse anti-H3K4me3 (Invitrogen). Secondary antibodies were purchased from Invitrogen. DNA was labeled with DAPI (Invitrogen). Embryos were collected and fixed via a formaldehyde/MeOH method [10]. Polytene chromosome preparations and staining were performed as in Karachentsev *et al*. [29]. Images of the ventral nerve cord were obtained using a Leica SP8 using a 40x Objective. Fas2 and HRP labeled embryos were imaged and typically contained 8-10 hemisegments. Hemisegments were examined for midline crossing and in some instances the presence or integrity of the most lateral Fas2+ longitudinal track. Similar imaging and analysis were performed on Connectin/HRP labeled embryos.

### LC-MS/MS for 5mC and 5hmC detection and quantification

Mass spectrometry analysis was performed as described previously [30]. Briefly, 3 μL of 10× buffer (500 mM Tris-HCl, 100 mM NaCl, 10 mM MgCl_2_, 10 mM ZnSO_4_, pH 7.0), 2 μL (180 units) of S1 nuclease, 2 μL (0.001 units) of venom phosphodiesterase I and 1 μL (30 units) of CAIP were added to 1 μg of mRNA from *Drosophila* wild type and Tet-deficient larval brains, respectively (in 22 μL of H_2_O). The mixture (30 μL) was incubated at 37°C for 4 h. The resulting solution was three times extracted with chloroform. The upper aqueous phase was collected and passed through a solid phase extraction cartridge filled with 50 mg of sorbent of graphitized carbon black to remove the salts. The eluate was then dried with nitrogen at 37°C for subsequent chemical labeling and LC-ESI-MS/MS analysis by an AB 3200 QTRAP mass spectrometer (Applied Biosystems, Foster City, CA, USA).

### Embryo and Larval Tet ChIP-seq

0-12h embryos were collected, processed, and chromatin was prepared according to Yad *et al.* [31], except lysates were sonicated on a Covaris S2 sonication device (intensity 8, duty cycle 20%, cycle burst 200) for 30 minutes at 4°C to reach fragments ranging from 150–500 bp and then centrifuged at 20,000g at 4°C for 1 minute. Supernatants were collected and centrifuged again for 15 minute to remove debris. Chromatin samples were then snap frozen in dry ice and stored at -80°C until immunoprecipitation in triplicates. All buffers contained cOmplete EDTA-free protease inhibitor cocktail (Roche).

For the larval brain fraction (LBF), 300 frozen larval heads were thawed on ice and 1 ml of NU-1 buffer (5 mM HEPES-KOH pH 7.9, 5 mM MgCl_2_, 0.1 mM EDTA pH 8.0, 0.5 mM EGTA pH 8.0, 350 mM sucrose, 1mM DTT). 1% formaldehyde was added to NU-1 buffer before use. Samples were homogenized immediately at room temperature using Dounce with a loose pestle 30 times without foaming for 15 minutes. Samples were filtered first through BD Falcon Cell Strainer 70 μm (Cat No.352350) followed by 50 μm Falcon (Cat No. 340603). Samples were quenched with freshly prepared 125 mM glycine incubated for 5 minutes at room temperature on a shaker and transferred to ice for 5 minutes. Samples were centrifuged at 4000 g at 4°C for 5 minutes. The pellet was washed twice with 1 ml cold PBS and resuspended in 350 µl chilled sonication buffer (50mM HEPES-KOH pH 7.9, 140 mM NaCl, 1mM EDTA pH 8.0, 1% Triton X-100, 0.1% sodium deoxycholate, 1% SDS) and incubated for 20 minutes at 4°C. Lysates were sonicated as described above and chromatin was stored at -80°C until immunoprecipitation.

### Chromatin Immunoprecipitation

Chromatin samples were thawed on ice and pre-cleared for 15 minutes by rotation in 25 µl of pre-washed binding control magnetic agarose beads (Chromotek). Chromatin was diluted ten-fold in sonication buffer without SDS. 1% of the diluted lysate was recovered and used as input. Diluted chromatin was incubated with 25 µl of pre-washed GFP-Trap MA beads (Chromotek) and rotated at 4°C overnight. Lysates were washed on magnetic stand with 1 ml each low salt RIPA buffer (140 mM NaCl, 1mM EDTA pH 8.0, 1% Triton X-100, 0.1% sodium deoxycholate, 10mM Tris-HCl pH 8.0) (5 times), high salt RIPA buffer (500 mM NaCl, 1mM EDTA pH 8.0, 1% Triton X-100, 0.1% sodium deoxycholate, 10mM Tris-HCl pH 8.0) (2 times), LiCl buffer (250mM LiCl, 1mM EDTA pH 8.0, 0.5% IGEPAL CA-630, 0.5% sodium deoxycholate, 10mM Tris-HCl pH 8.0) (1 time), TE buffer (10mM Tris-HCl pH 8.0, 1mM EDTA pH 8.0) (1 time). All buffers contained cOmplete EDTA-free protease inhibitor cocktail (Roche).

ChIP DNA was eluted by shaking 2 hours at 37°C with 100 µl of elution buffer (1% SDS, 50mM NaHCO_3_, 10µg/ml RNaseA), then 4 hours with 0.2µg/ml proteinase K. Beads were concentrated on magnet and elute was recovered. Samples were de-crosslinked overnight at 65°C. Inputs were processed like ChIP samples. DNA was purified by phenol/chloroform/isoamyl alcohol followed by SPRI select beads (Beckman Coulter) and DNA concentration was measured with Qubit fluorometer (Thermo Fisher).

### Embryo Tet ChIP-seq library preparation and sequencing

NGS Libraries were made from eluted DNA using the NEBNext Ultra II DNA Library Prep kit (New England Biolabs) according to the manufacturer’s protocol. Briefly, 20 ng of DNA fragments were end-repaired and the blunt, phosphorylated ends were treated with Klenow DNA polymerase and dATP to yield a 3′ A base overhang for ligation of Illumina adapters. After adapter ligation, DNA was PCR amplified with indexed primer for 12 cycles. Libraries were size-selected using Ampure XP beads (Beckman Coulter) to remove adapter dimers. DNA was quantified by fluorometry with the Qubit 2.0 (Thermo Scientific) and DNA integrity was assessed with a Fragment Analyzer (Agilent). The libraries were pooled and sequenced on the NextSeq 500 platform using 75 bp single end sequencing according to manufacturer’s protocol using Reagent v.2.5 at the Waksman Institute Genomics Core. Coverage ranged from 30 million to 60 million tags per ChIP-seq sample.

### Larva Tet ChIP-seq library preparation and sequencing

ACCEL-NGS® 1S plus DNA library kit was used to prepare indexed libraries from IP and input DNA. Libraries were pooled respecting equimolarity. Sequencing was performed on Illumina MISeq sequencer in 150 bp paired-end reads.

### Embryo Tet ChIP-seq data analysis

Raw reads were trimmed using cutadapt v2.0 [32] to remove adapter and low-quality reads. The processed reads were mapped to the *Drosophila melanogaster* BDGP6 (dm6) reference genome from Ensembl release 88 using the BWA version 0.7.5-r404 for Chip-seq [33]. For analysis, only unique reads with mapping quality >20 were accepted. Further, redundant reads with identical coordinates were filtered out. Aligned reads were processed by Model-based Analysis of ChIP-seq (MACS2) [34] using Input ChIP DNA as control. For peak calling the MACS2 ‘callpeak’ function was used (-p 1e-2 -g 1.2e+08 -B --nomodel –ext size 147 –SPMR) for each replicate vs. control input. Peaks were selected using the following criteria: p-value <10e-5, fold enrichment over control greater than 10 and a minimal number of reads higher than 50. Bedtools (version v2.24.0) [35] was used to identify overlapping peaks in replicates. A sliding window of 50, 100, 150, 200, 250 and 300 bp around the peak summit (base position of maximum enrichment) was used to determine best range for overlapping peaks. The number of overlapping peaks saturated around window size of 250 bp. Thus, for downstream analysis, windows size of 250 bp was used to identify overlapping peaks in replicates. The Integrated Genomics Viewer (IGV) [36] was used for visualization of ChIP-seq data sets. For visualization in IGV, bigwig peak files were generated using “bdgcmp” function in MACS2 with option “-m logFE -p 0.00001”. Peaks were annotated using the “annotatePeaks.pl” feature of HomerTools [37] with default settings and gtf was obtained from of Ensembl dm6 release 88. De novo motif discovery was carried out on all intersecting peaks of Tet ChIP-seq. DNA sequences (FASTA) were generated from chromosome coordinates produced by peak detection and windowing using the BEDTools. De novo motif analysis was performed using MEME-ChIP [38]. Gene ontology (GO) analysis was done using Database for Annotation, Visualization and Integrated Discovery (DAVID) [39, 40]. Binding profile within gene body was generated using deepTools2 with computeMatrix and plotProfile functions [41].

### H3K4me3 ChIP-seq public datasets and analysis

Embryo and larva H3K4me3 ChIP-seq data were obtained from the modENCODE project (GEO: GSE16013) [42]. The analysis was carried out from raw data following the same approach described for Tet ChIP-seq. The overlapping of Tet-ChIP seq peaks and H3K4me3 was computed using BEDTools [35].

### Larva Tet ChIP-seq data analysis

Tet-Chip sequencing data were pre-processed using the following steps: the raw sequencing data were first analysed with FastQC (Andrews, 2010, https://www.bioinformatics.babraham.ac.uk/projects/fastqc/). Low-complexity reads were removed with the AfterQC tool [43] with default parameters and Trimmomatic [44] with default parameters was used to remove adapter sequences. The resulting fastq data were again analysed with FastQC to ensure that no further processing was needed. Pre-processed reads were then mapped against the *Drosophila* reference genome (BDGP6.28) with the bowtie2 algorithm [45] using the ensembl reference transcriptome (version 100). Tet-binding peak regions were identified by applying the MACS2 peak-calling tool [34] to immunoprecipitated (IP) samples, using their input counterpart to estimate background noise (q-value < 0.05). It is worth noting that the “expected genome size” MACS2 parameter was set as the *Drosophila* genome length excluding ‘N’ bases (*i.e.,* 142 573 024 bp), and summit positions were identified using the MACS2 “-call-summits” option. To avoid identifying extremely large peak regions, the peaks were resized to 100 bp on both sides of the identified summit. Binding profile within gene body was generated using deepTools2 with computeMatrix and plotProfile functions [41].

### HydroxyMethylated RNA Immunoprecipitation sequencing (hMeRIP-seq)

0-12h embryos were collected, immediately frozen on dry ice, and stored at -80°C until RNA purification. The larval brain fraction (LBF), was dissected, immediately frozen on dry ice, and stored at -80°C until RNA isolation. The RNA immunoprecipitation was performed essentially as described in Dominissi *et al.* [46]. Briefly, total RNA was isolated using RNeasy Maxi Kit (Qiagen). For each sample 1 mg of total RNA (1 μg/μl) was divided into batches of 45µg and incubated at 94°C in fragmentation buffer (100 mM Tris-HCl pH7.0, 100 mM ZnCl_2_) for 40 seconds. Fragmented RNA batches were pooled, and ethanol precipitated at -80°C overnight. RNA samples were washed with 75% ethanol and resuspended in RNase-free water. Fragmentation efficiency was checked on a Bioanalyzer RNA chip (Agilent). RNA fragments were denatured by heating at 70°C for 5 minutes, then chilled on ice for 5 minutes. For immunoprecipitation, RNA samples were incubated overnight at 4°C with 12.5 µg of anti-5-hmC antibody (Diagenode rat monoclonal MAb-633HMC) or without antibody as negative control in IP buffer (750 mM NaCl, 50 mM Tris-HCl pH7.4, 0.5% IGEPAL CA-630, RNasin 400 U/ml and RVC 2 mM). 60 μl of equilibrated Dynabeads Protein G (Life Technologies) were added to the samples and incubated at 4°C for 2.5 hours. The magnetic stand beads were washed with 1 ml IP buffer for 5 minutes three times. To elute immunoprecipitated RNA, 1 ml TriPure Reagent (Roche) was added, mixed thoroughly, and centrifuged at room temperature for 5 minutes. Aqueous phase was recovered, and equal amount of chloroform was added, vortexed and aqueous phase was collected after centrifugation and ethanol precipitated at -80°C overnight. RNA was resuspended in nuclease free water and used for library preparation. All buffers contained cOmplete EDTA-free protease inhibitor cocktail (Roche).

### hMeRIP-seq library preparation and sequencing

Library preparation was done with the TruSeq ChIP Sample Prep Kit (Illumina) after reverse transcription of pulled-down RNA and synthesis of a second strand (NEB) by Next mRNA second strand synthesis module (NEB)). Briefly, 5 to 10 ng dsDNA was subjected to 5’ and 3’ protruding end repair. Then, non-templated adenines were added to the 3’ ends of the blunted DNA fragments. This last step allows ligation of Illumina multiplex adapters. The DNA fragments were then size selected in order to remove all unligated adapters and to sequence 200-300-bp fragments. 18 cycles of PCR were carried out to amplify the library. DNA was quantified by fluorometry with the Qubit 2.0 and DNA integrity was assessed with a 2100 bioanalyzer (Agilent). 6 pM of DNA library spiked with .5% PhiX viral DNA was clustered on cBot (Illumina) and then sequenced on a HiScanSQ module (Illumina).

### hMeRIP-seq data analysis

The processed reads were mapped to the reference genome Drosophila melanogaster BDGP6 (dm6) from Ensembl by using Hisat2 (version 2.1.0) for RNA seq and hMeRIP seq [47]. To analyze gene expression, HTSeq framework, version 0.5.3p9, was used to count the aligned reads in genes [48]. Mode “union” and mapping quality cut-off 20 were used for our analysis. Count-table was normalized so that all samples have the same level of total mapped reads. DEseq2 was used to identify differentially expressed genes [49]. Cufflinks v2.2.1 was applied to calculate the rpkm values [50, 51]. A gene was considered as significantly changed when fold change >=2 or <= -2 and adjusted p value < 0.05. “SplitNCigarReads” funciton in GATK (version 3.3-0) (https://gatk.broadinstitute.org/) were used to split reads that contain Ns in their cigar string (e.g., spanning splicing events in hMeRIP-seq data). “rmdup” function of samtools (version 1.3.1) were used to remove a duplicate mapping of reads. Then the same peak calling procedure as ChIP seq data analysis was performed to call peaks of hMeRIP-seq data. The peaks of hMeRIP-seq were selected using P-value < 10e-5. Peaks of hMeRIP-seq were considered as reduced when the normalized hMeRIP-seq signal in control samples was at least 1.4-fold change higher than the signal in Tet depleted samples. The fold change and P-value were calculated using “limma” package in R [52].

### Western blot

One hundred third instar larval brains from wild type or *Tet^null^*were dissected and immediately frozen on dry ice. Total protein was isolated from these brains using RIPA buffer and 75 ug of the total protein was loaded to each well. Slit antibody (DSHB, C555.6D, Spyros Artavanis-Tsakonas) was used at 1: 200 dilution and Robo2 antibody [53] was used at 1: 1000 dilution. The western blot signals were detected using IRDye 800CW Infrared Dyes conjugated secondary antibody in LICOR Odyssey CLx imaging system. Signals were quantified using LICOR Image Studio Lite software. See Figure S8 for unprocessed western blot exposure.

### Statistical information

Statistical analysis was performed using R or GraphPad Prism 9. Statistics were performed using Student’s t-test or chi-square test unless otherwise specified. Error bars are presented as SEM. P-value < 0.05 is the cut-off for statistical significance.

## Data availability

The sequencing data that support the finding of this study are available at NCBI Gene Expression Omnibus (GEO) with the accession number GSE225980 and accessible token “ihmzwuocfrybdal”.

## Acknowledgements

We thank Cordelia Rauskolb, Bryce Nickels, and Michael Verzi for helpful comments on the manuscript, Premal Shah, John Favate, and Shun Liang for discussions and suggestions, and Benjamin Rogers-Boehme for help with Figures. We also thank Barry Dickson for anti-Robo2 antibodies and Le Nguyen for expert fly food preparation and stock maintenance. Stocks obtained from the Bloomington Drosophila Stock Center (NIHP40OD018537) were used in this study. The imaging was done at the Waksman Institute Shared Imaging Facility, Rutgers University.

## References

1. Roundtree IA, Evans ME, Pan T, He C. Dynamic RNA Modifications in Gene Expression Regulation. Cell. 2017;169(7):1187–200. doi: 10.1016/j.cell.2017.05.045. PubMed PMID: 28622506; PubMed Central PMCID: PMCPMC5657247.

2. Boccaletto P, Stefaniak F, Ray A, Cappannini A, Mukherjee S, Purta E, et al. MODOMICS: a database of RNA modification pathways. 2021 update. Nucleic Acids Res. 2022;50(D1):D231–D5. doi: 10.1093/nar/gkab1083. PubMed PMID: 34893873; PubMed Central PMCID: PMCPMC8728126.

3. Schaefer MR. The Regulation of RNA Modification Systems: The Next Frontier in Epitranscriptomics? Genes (Basel). 2021;12(3). Epub 20210226. doi: 10.3390/genes12030345. PubMed PMID: 33652758; PubMed Central PMCID: PMCPMC7996938.

4. Gao Y, Fang J. RNA 5-methylcytosine modification and its emerging role as an epitranscriptomic mark. RNA Biol. 2021;18(sup1):117–27. Epub 20210721. doi: 10.1080/15476286.2021.1950993. PubMed PMID: 34288807; PubMed Central PMCID: PMCPMC8677007.

5. Boffelli D, Takayama S, Martin DI. Now you see it: genome methylation makes a comeback in Drosophila. Bioessays. 2014;36(12):1138–44. Epub 20140912. doi: 10.1002/bies.201400097. PubMed PMID: 25220261.

6. Delatte B, Wang F, Ngoc LV, Collignon E, Bonvin E, Deplus R, et al. RNA biochemistry. Transcriptome-wide distribution and function of RNA hydroxymethylcytosine. Science. 2016;351(6270):282–5. Epub 2016/01/28. doi: 10.1126/science.aac5253. PubMed PMID: 26816380.

7. Tahiliani M, Koh KP, Shen Y, Pastor WA, Bandukwala H, Brudno Y, et al. Conversion of 5-methylcytosine to 5-hydroxymethylcytosine in mammalian DNA by MLL partner TET1. Science. 2009;324(5929):930–5. Epub 20090416. doi: 10.1126/science.1170116. PubMed PMID: 19372391; PubMed Central PMCID: PMCPMC2715015.

8. Fu L, Guerrero CR, Zhong N, Amato NJ, Liu Y, Liu S, et al. Tet-mediated formation of 5-hydroxymethylcytosine in RNA. J Am Chem Soc. 2014;136(33):11582–5. Epub 20140807. doi: 10.1021/ja505305z. PubMed PMID: 25073028; PubMed Central PMCID: PMCPMC4140497.

9. Tan L, Shi YG. Tet family proteins and 5-hydroxymethylcytosine in development and disease. Development. 2012;139(11):1895–902. doi: 10.1242/dev.070771. PubMed PMID: 22569552; PubMed Central PMCID: PMCPMC3347683.

10. Wang F, Minakhina S, Tran H, Changela N, Kramer J, Steward R. Tet protein function during Drosophila development. PLoS One. 2018;13(1):e0190367. Epub 20180111. doi: 10.1371/journal.pone.0190367. PubMed PMID: 29324752; PubMed Central PMCID: PMCPMC5764297.

11. Dunwell TL, McGuffin LJ, Dunwell JM, Pfeifer GP. The mysterious presence of a 5-methylcytosine oxidase in the Drosophila genome: possible explanations. Cell Cycle. 2013;12(21):3357–65. Epub 20130919. doi: 10.4161/cc.26540. PubMed PMID: 24091536; PubMed Central PMCID: PMCPMC3895424.

12. Zhang G, Huang H, Liu D, Cheng Y, Liu X, Zhang W, et al. N6-methyladenine DNA modification in Drosophila. Cell. 2015;161(4):893–906. Epub 20150430. doi: 10.1016/j.cell.2015.04.018. PubMed PMID: 25936838.

13. Ismail JN, Badini S, Frey F, Abou-Kheir W, Shirinian M. Drosophila Tet Is Expressed in Midline Glia and Is Required for Proper Axonal Development. Front Cell Neurosci. 2019;13:252. Epub 20190604. doi: 10.3389/fncel.2019.00252. PubMed PMID: 31213988; PubMed Central PMCID: PMCPMC6558204.

14. Shen Q, Zhang Q, Shi Y, Shi Q, Jiang Y, Gu Y, et al. Tet2 promotes pathogen infection-induced myelopoiesis through mRNA oxidation. Nature. 2018;554(7690):123–7. Epub 20180124. doi: 10.1038/nature25434. PubMed PMID: 29364877.

15. Guallar D, Bi X, Pardavila JA, Huang X, Saenz C, Shi X, et al. RNA-dependent chromatin targeting of TET2 for endogenous retrovirus control in pluripotent stem cells. Nat Genet. 2018;50(3):443–51. Epub 20180226. doi: 10.1038/s41588-018-0060-9. PubMed PMID: 29483655; PubMed Central PMCID: PMCPMC5862756.

16. Lan J, Rajan N, Bizet M, Penning A, Singh NK, Guallar D, et al. Functional role of Tet-mediated RNA hydroxymethylcytosine in mouse ES cells and during differentiation. Nat Commun. 2020;11(1):4956. Epub 20201002. doi: 10.1038/s41467-020-18729-6. PubMed PMID: 33009383; PubMed Central PMCID: PMCPMC7532169.

17. Wu H, D’Alessio AC, Ito S, Xia K, Wang Z, Cui K, et al. Dual functions of Tet1 in transcriptional regulation in mouse embryonic stem cells. Nature. 2011;473(7347):389–93. Epub 20110330. doi: 10.1038/nature09934. PubMed PMID: 21451524; PubMed Central PMCID: PMCPMC3539771.

18. Jones PA, Liang G. Rethinking how DNA methylation patterns are maintained. Nat Rev Genet. 2009;10(11):805–11. Epub 20090930. doi: 10.1038/nrg2651. PubMed PMID: 19789556; PubMed Central PMCID: PMCPMC2848124.

19. Blockus H, Chedotal A. Slit-Robo signaling. Development. 2016;143(17):3037–44. doi: 10.1242/dev.132829. PubMed PMID: 27578174.

20. Simpson JH, Kidd T, Bland KS, Goodman CS. Short-range and long-range guidance by slit and its Robo receptors. Robo and Robo2 play distinct roles in midline guidance. Neuron. 2000;28(3):753–66. doi: 10.1016/s0896-6273(00)00151-3. PubMed PMID: 11163264.

21. Evans TA, Santiago C, Arbeille E, Bashaw GJ. Robo2 acts in trans to inhibit Slit-Robo1 repulsion in pre-crossing commissural axons. Elife. 2015;4:e08407. Epub 20150717. doi: 10.7554/eLife.08407. PubMed PMID: 26186094; PubMed Central PMCID: PMCPMC4505356.

22. Beck DB, Petracovici A, He C, Moore HW, Louie RJ, Ansar M, et al. Delineation of a Human Mendelian Disorder of the DNA Demethylation Machinery: TET3 Deficiency. Am J Hum Genet. 2020;106(2):234–45. Epub 20200109. doi: 10.1016/j.ajhg.2019.12.007. PubMed PMID: 31928709; PubMed Central PMCID: PMCPMC7010978.

23. Gorla M, Bashaw GJ. Molecular mechanisms regulating axon responsiveness at the midline. Dev Biol. 2020;466(1-2):12–21. Epub 20200817. doi: 10.1016/j.ydbio.2020.08.006. PubMed PMID: 32818516; PubMed Central PMCID: PMCPMC8447865.

24. Dominissini D, Moshitch-Moshkovitz S, Schwartz S, Salmon-Divon M, Ungar L, Osenberg S, et al. Topology of the human and mouse m6A RNA methylomes revealed by m6A-seq. Nature. 2012;485(7397):201–6. Epub 20120429. doi: 10.1038/nature11112. PubMed PMID: 22575960.

25. Yao B, Li Y, Wang Z, Chen L, Poidevin M, Zhang C, et al. Active N(6)-Methyladenine Demethylation by DMAD Regulates Gene Expression by Coordinating with Polycomb Protein in Neurons. Mol Cell. 2018;71(5):848–57 e6. Epub 20180802. doi: 10.1016/j.molcel.2018.07.005. PubMed PMID: 30078725; PubMed Central PMCID: PMCPMC6136845.

26. Simpson JH, Bland KS, Fetter RD, Goodman CS. Short-range and long-range guidance by Slit and its Robo receptors: a combinatorial code of Robo receptors controls lateral position. Cell. 2000;103(7):1019–32. doi: 10.1016/s0092-8674(00)00206-3. PubMed PMID: 11163179.

27. Spitzweck B, Brankatschk M, Dickson BJ. Distinct protein domains and expression patterns confer divergent axon guidance functions for Drosophila Robo receptors. Cell. 2010;140(3):409–20. doi: 10.1016/j.cell.2010.01.002. PubMed PMID: 20144763.

28. Santiago-Martinez E, Soplop NH, Kramer SG. Lateral positioning at the dorsal midline: Slit and Roundabout receptors guide Drosophila heart cell migration. Proc Natl Acad Sci U S A. 2006;103(33):12441–6. Epub 20060803. doi: 10.1073/pnas.0605284103. PubMed PMID: 16888037; PubMed Central PMCID: PMCPMC1567898.

29. Karachentsev D, Sarma K, Reinberg D, Steward R. PR-Set7-dependent methylation of histone H4 Lys 20 functions in repression of gene expression and is essential for mitosis. Genes Dev. 2005;19(4):431–5. Epub 20050128. doi: 10.1101/gad.1263005. PubMed PMID: 15681608; PubMed Central PMCID: PMCPMC548943.

30. Huang W, Lan MD, Qi CB, Zheng SJ, Wei SZ, Yuan BF, et al. Formation and determination of the oxidation products of 5-methylcytosine in RNA. Chem Sci. 2016;7(8):5495–502. Epub 20160511. doi: 10.1039/c6sc01589a. PubMed PMID: 30034689; PubMed Central PMCID: PMCPMC6021781.

31. Ghavi-Helm Y, Zhao B, Furlong EE. Chromatin Immunoprecipitation for Analyzing Transcription Factor Binding and Histone Modifications in Drosophila. Methods Mol Biol. 2016;1478:263–77. doi: 10.1007/978-1-4939-6371-3_16. PubMed PMID: 27730588.

32. Martin M. Cutadapt removes adapter sequences from high-throughput sequencing reads. 2011. 2011;17(1):3. Epub 2011-08-02. doi: 10.14806/ej.17.1.200.

33. Li H, Durbin R. Fast and accurate short read alignment with Burrows-Wheeler transform. Bioinformatics. 2009;25(14):1754–60. Epub 20090518. doi: 10.1093/bioinformatics/btp324. PubMed PMID: 19451168; PubMed Central PMCID: PMCPMC2705234.

34. Zhang Y, Liu T, Meyer CA, Eeckhoute J, Johnson DS, Bernstein BE, et al. Model-based analysis of ChIP-Seq (MACS). Genome Biol. 2008;9(9):R137. Epub 20080917. doi: 10.1186/gb-2008-9-9-r137. PubMed PMID: 18798982; PubMed Central PMCID: PMCPMC2592715.

35. Quinlan AR, Hall IM. BEDTools: a flexible suite of utilities for comparing genomic features. Bioinformatics. 2010;26(6):841–2. Epub 20100128. doi: 10.1093/bioinformatics/btq033. PubMed PMID: 20110278; PubMed Central PMCID: PMCPMC2832824.

36. Robinson JT, Thorvaldsdottir H, Winckler W, Guttman M, Lander ES, Getz G, et al. Integrative genomics viewer. Nat Biotechnol. 2011;29(1):24–6. doi: 10.1038/nbt.1754. PubMed PMID: 21221095; PubMed Central PMCID: PMCPMC3346182.

37. Heinz S, Benner C, Spann N, Bertolino E, Lin YC, Laslo P, et al. Simple combinations of lineage-determining transcription factors prime cis-regulatory elements required for macrophage and B cell identities. Mol Cell. 2010;38(4):576–89. doi: 10.1016/j.molcel.2010.05.004. PubMed PMID: 20513432; PubMed Central PMCID: PMCPMC2898526.

38. Machanick P, Bailey TL. MEME-ChIP: motif analysis of large DNA datasets. Bioinformatics. 2011;27(12):1696–7. Epub 20110412. doi: 10.1093/bioinformatics/btr189. PubMed PMID: 21486936; PubMed Central PMCID: PMCPMC3106185.

39. Huang da W, Sherman BT, Lempicki RA. Systematic and integrative analysis of large gene lists using DAVID bioinformatics resources. Nat Protoc. 2009;4(1):44–57. doi: 10.1038/nprot.2008.211. PubMed PMID: 19131956.

40. Sherman BT, Hao M, Qiu J, Jiao X, Baseler MW, Lane HC, et al. DAVID: a web server for functional enrichment analysis and functional annotation of gene lists (2021 update). Nucleic Acids Res. 2022;50(W1):W216–21. Epub 20220323. doi: 10.1093/nar/gkac194. PubMed PMID: 35325185; PubMed Central PMCID: PMCPMC9252805.

41. Ramirez F, Ryan DP, Gruning B, Bhardwaj V, Kilpert F, Richter AS, et al. deepTools2: a next generation web server for deep-sequencing data analysis. Nucleic Acids Res. 2016;44(W1):W160–5. Epub 20160413. doi: 10.1093/nar/gkw257. PubMed PMID: 27079975; PubMed Central PMCID: PMCPMC4987876.

42. Negre N, Brown CD, Ma L, Bristow CA, Miller SW, Wagner U, et al. A cis-regulatory map of the Drosophila genome. Nature. 2011;471(7339):527–31. doi: 10.1038/nature09990. PubMed PMID: 21430782; PubMed Central PMCID: PMCPMC3179250.

43. Chen S, Huang T, Zhou Y, Han Y, Xu M, Gu J. AfterQC: automatic filtering, trimming, error removing and quality control for fastq data. BMC Bioinformatics. 2017;18(Suppl 3):80. Epub 20170314. doi: 10.1186/s12859-017-1469-3. PubMed PMID: 28361673; PubMed Central PMCID: PMCPMC5374548.

44. Bolger AM, Lohse M, Usadel B. Trimmomatic: a flexible trimmer for Illumina sequence data. Bioinformatics. 2014;30(15):2114–20. Epub 20140401. doi: 10.1093/bioinformatics/btu170. PubMed PMID: 24695404; PubMed Central PMCID: PMCPMC4103590.

45. Langmead B, Salzberg SL. Fast gapped-read alignment with Bowtie 2. Nat Methods. 2012;9(4):357–9. Epub 20120304. doi: 10.1038/nmeth.1923. PubMed PMID: 22388286; PubMed Central PMCID: PMCPMC3322381.

46. Dominissini D, Moshitch-Moshkovitz S, Salmon-Divon M, Amariglio N, Rechavi G. Transcriptome-wide mapping of N(6)-methyladenosine by m(6)A-seq based on immunocapturing and massively parallel sequencing. Nat Protoc. 2013;8(1):176–89. Epub 20130103. doi: 10.1038/nprot.2012.148. PubMed PMID: 23288318.

47. Kim D, Paggi JM, Park C, Bennett C, Salzberg SL. Graph-based genome alignment and genotyping with HISAT2 and HISAT-genotype. Nat Biotechnol. 2019;37(8):907–15. Epub 20190802. doi: 10.1038/s41587-019-0201-4. PubMed PMID: 31375807; PubMed Central PMCID: PMCPMC7605509.

48. Anders S, Pyl PT, Huber W. HTSeq--a Python framework to work with high-throughput sequencing data. Bioinformatics. 2015;31(2):166–9. Epub 20140925. doi: 10.1093/bioinformatics/btu638. PubMed PMID: 25260700; PubMed Central PMCID: PMCPMC4287950.

49. Love MI, Huber W, Anders S. Moderated estimation of fold change and dispersion for RNA-seq data with DESeq2. Genome Biol. 2014;15(12):550. doi: 10.1186/s13059-014-0550-8. PubMed PMID: 25516281; PubMed Central PMCID: PMCPMC4302049.

50. Trapnell C, Williams BA, Pertea G, Mortazavi A, Kwan G, van Baren MJ, et al. Transcript assembly and quantification by RNA-Seq reveals unannotated transcripts and isoform switching during cell differentiation. Nat Biotechnol. 2010;28(5):511–5. Epub 20100502. doi: 10.1038/nbt.1621. PubMed PMID: 20436464; PubMed Central PMCID: PMCPMC3146043.

51. Trapnell C, Hendrickson DG, Sauvageau M, Goff L, Rinn JL, Pachter L. Differential analysis of gene regulation at transcript resolution with RNA-seq. Nat Biotechnol. 2013;31(1):46–53. Epub 20121209. doi: 10.1038/nbt.2450. PubMed PMID: 23222703; PubMed Central PMCID: PMCPMC3869392.

52. Ritchie ME, Phipson B, Wu D, Hu Y, Law CW, Shi W, et al. limma powers differential expression analyses for RNA-sequencing and microarray studies. Nucleic Acids Res. 2015;43(7):e47. Epub 20150120. doi: 10.1093/nar/gkv007. PubMed PMID: 25605792; PubMed Central PMCID: PMCPMC4402510.

53. Rajagopalan S, Vivancos V, Nicolas E, Dickson BJ. Selecting a longitudinal pathway: Robo receptors specify the lateral position of axons in the Drosophila CNS. Cell. 2000;103(7):1033–45. doi: 10.1016/s0092-8674(00)00207-5. PubMed PMID: 11163180.

