## Supporting information for "Tet-dependent 5-hydroxymethyl-Cytosine modification of mRNA regulates axon guidance genes in *Drosophila*"

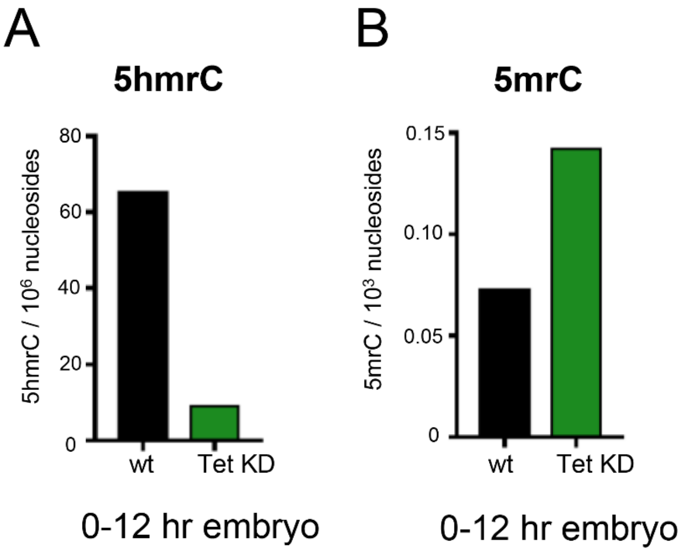


**S1 Fig. A. RNAi mediated KD of Tet alters the methylation status of Cytosine in total RNA by ultra-performance liquid chromatography tandem mass spectrometry.** 5hmrC in total RNA isolated from wild-type and Tet KD embryos. **B.** 5mrC in total RNA isolated from wild-type and Tet KD embryos.

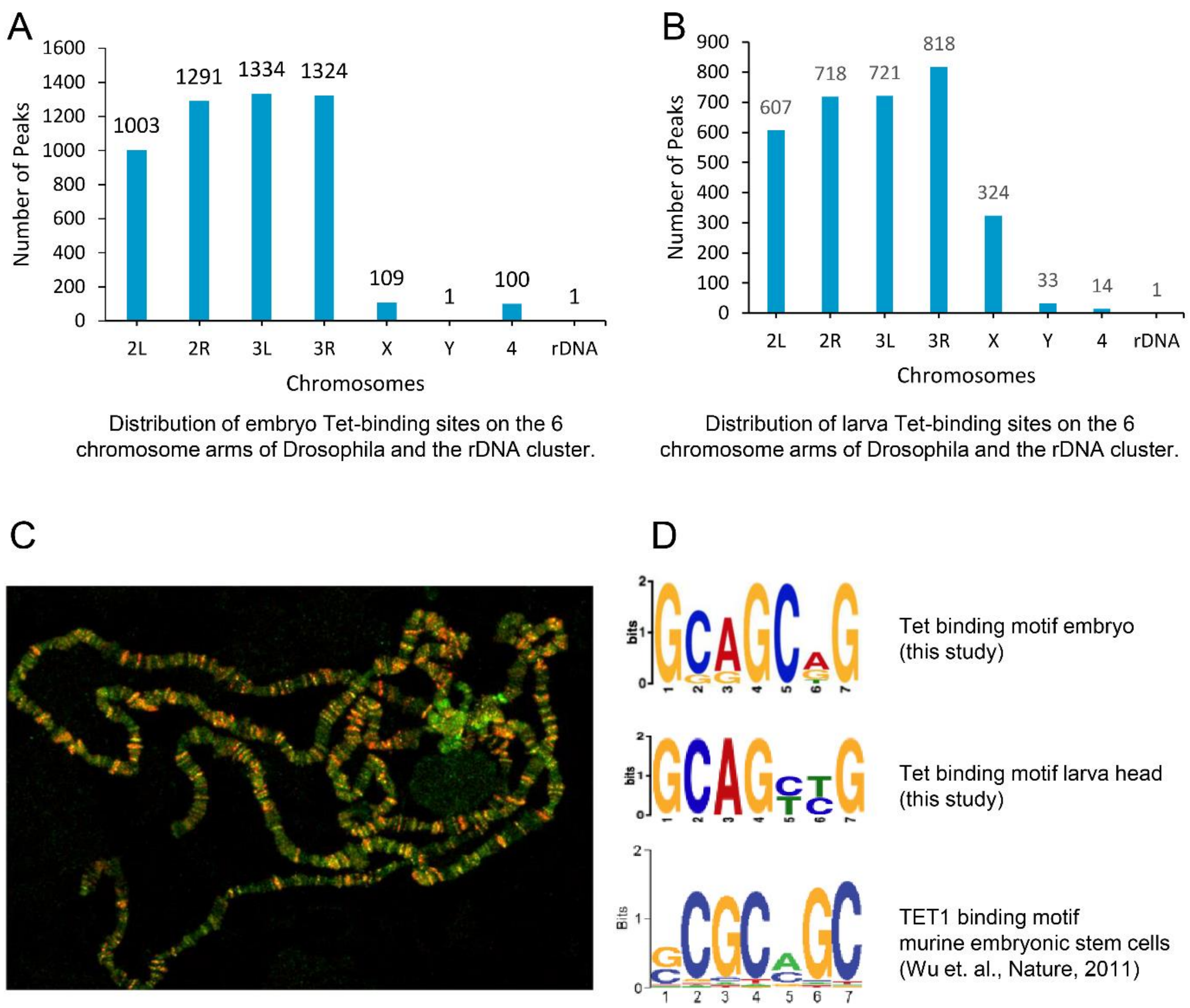


**S2 Fig. Chromsomal Distribution of Tet-binding. A**. Distribution of Tet-binding sites in embryo DNA on the 6 chromosome arms of Drosophila and the rDNA cluster (177 peaks are found at the centromeres or ummapped_scaffolds); **B.** Distribution of Tet-binding sites in LBF DNA. on the 6 chromosome arms of Drosophila and the rDNA cluster (17 peaks are found at the centromeres or ummapped scaffolds); **C.** Localization of Tet CxxC DNA-binding domain (red) on polythene chromosome. Salivary gland chromosome from hsp70-GAL4::UAS-TetCxxC-RFP-Myc 3rd instar larvae were stained with anti-Myc (red, TetCxxC) and H3K4me3 (green). Control chromosomes from *hsp70-GAL4* and UAS-TetCxxC alone show no Myc or RFP staining (not shown); **D.** Comparison of top DNA binding motives determined by Tet ChIP-seq in Drosophila 0-12 hr embryos, LBF, and murine ESCs**.**

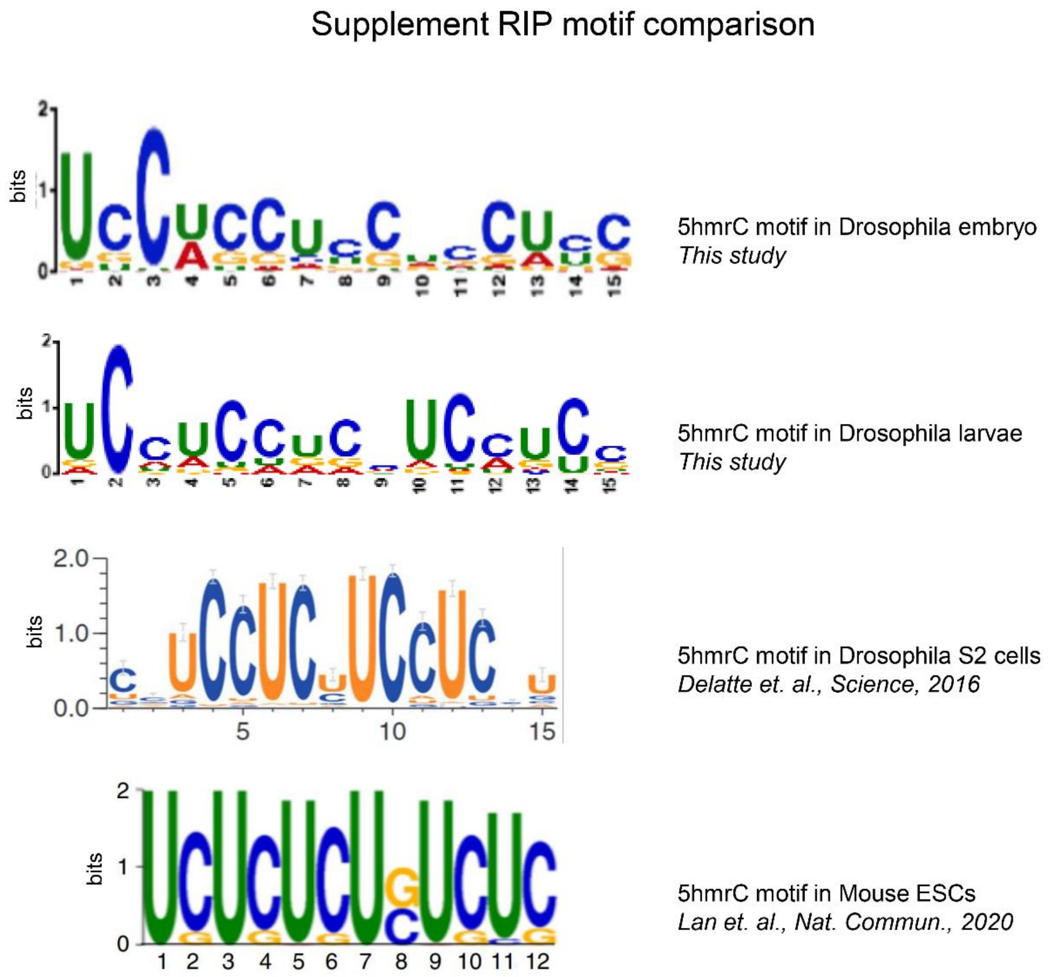


**S3 Fig**. **A. Comparison of the top sequence motives identified from 5hmrC peaks from Drosophila embryos.** LBF, Drosophila S2 cells and mouse ESCs.

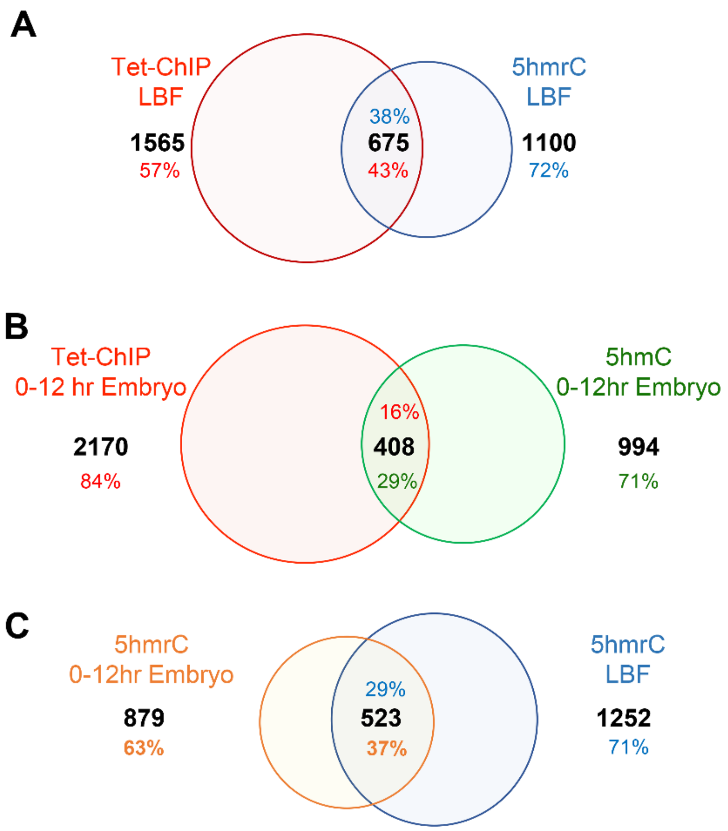


**S4 Fig. Overlap of Tet DNA binding genes with 5hmrC modified transcripts**. **A.,** in LBF, **B.**, in embryos. **C.** Overlap of 5hmrC modified transcripts in LBF and embryos.

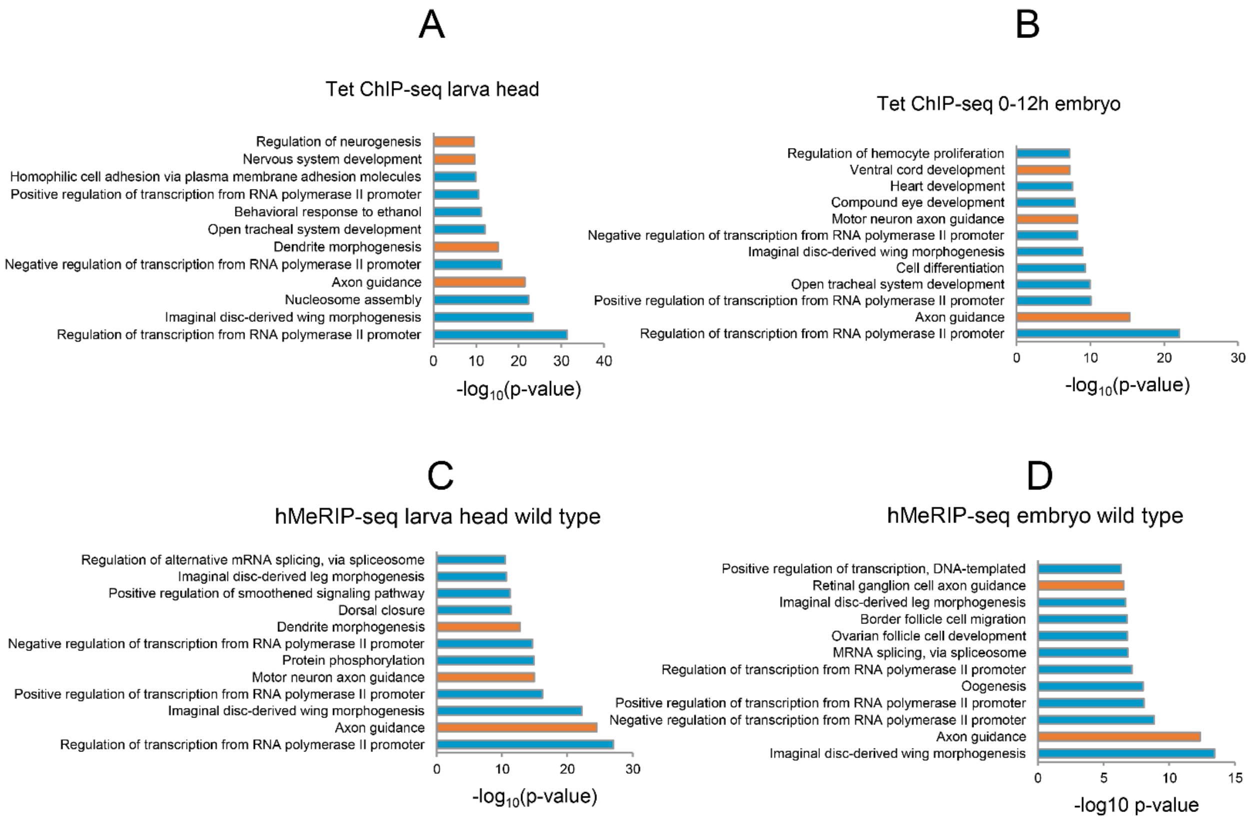


**S5 Fig**. **Comparison of GO term analyses. A.** of genome-wide Tet peaks from LBF; **B.** of genome-wide Tet peaks from 0-12 hour embryos; **C.** of transcriptome-wide 5hmrC peaks in wild type LBF; **D.** transcriptome-wide 5hmrC peaks in *Tet*$^{null}$ LBF. Note the consistency of the top two classes of genes.

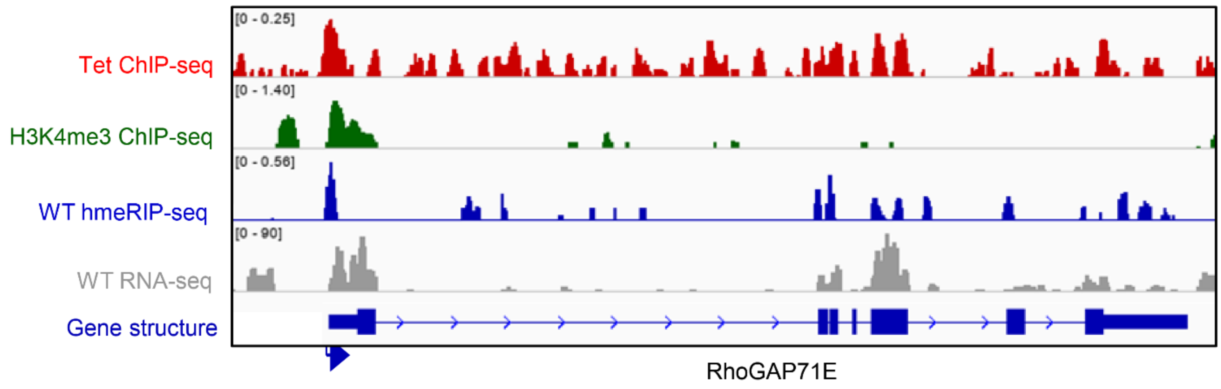


**S6 Fig.** Representative IGV tracks of RhoGAP71E gene showing Tet binding in 0-12 hr embryos. Y axis scale is indicated above each track.

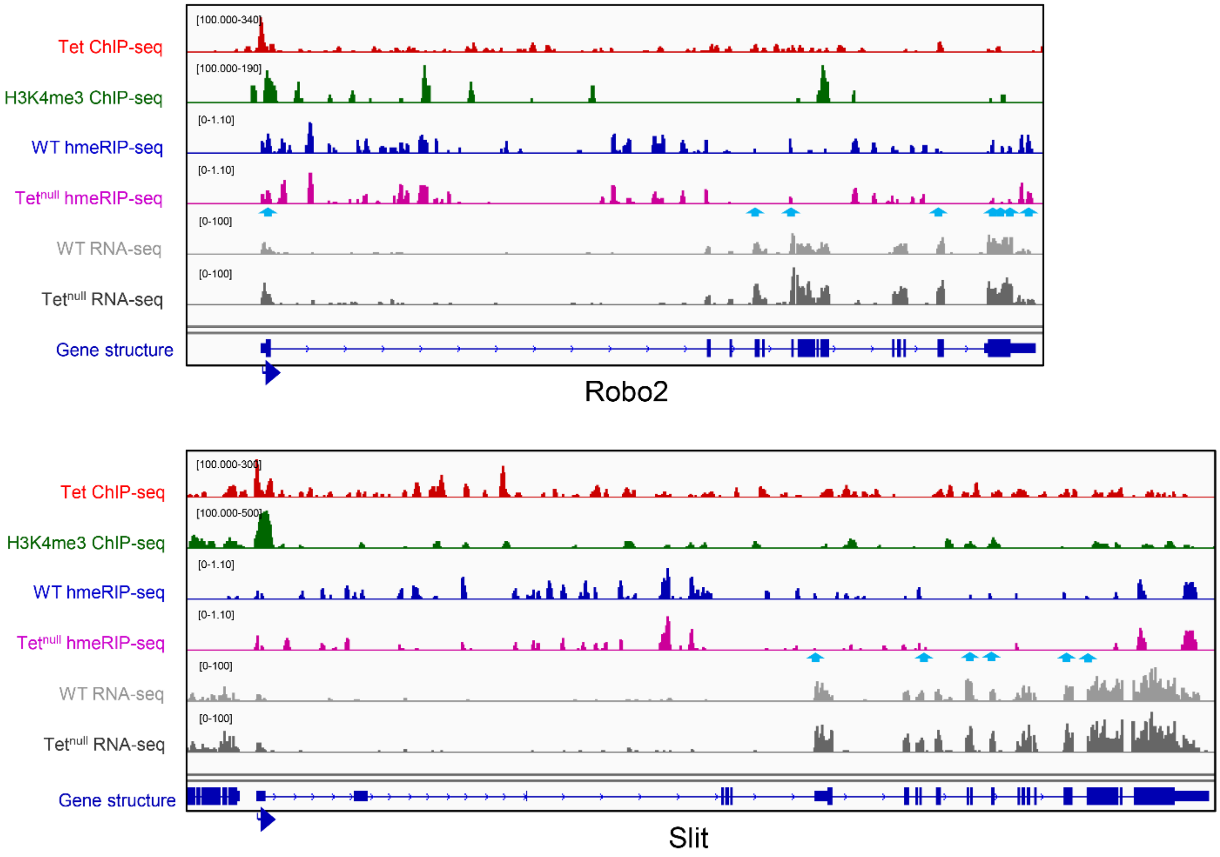


**S7 Fig. IGV tracks of Slit and Robo2 and Slit.** Blue arrows show reduction on 5hmrC peaks.

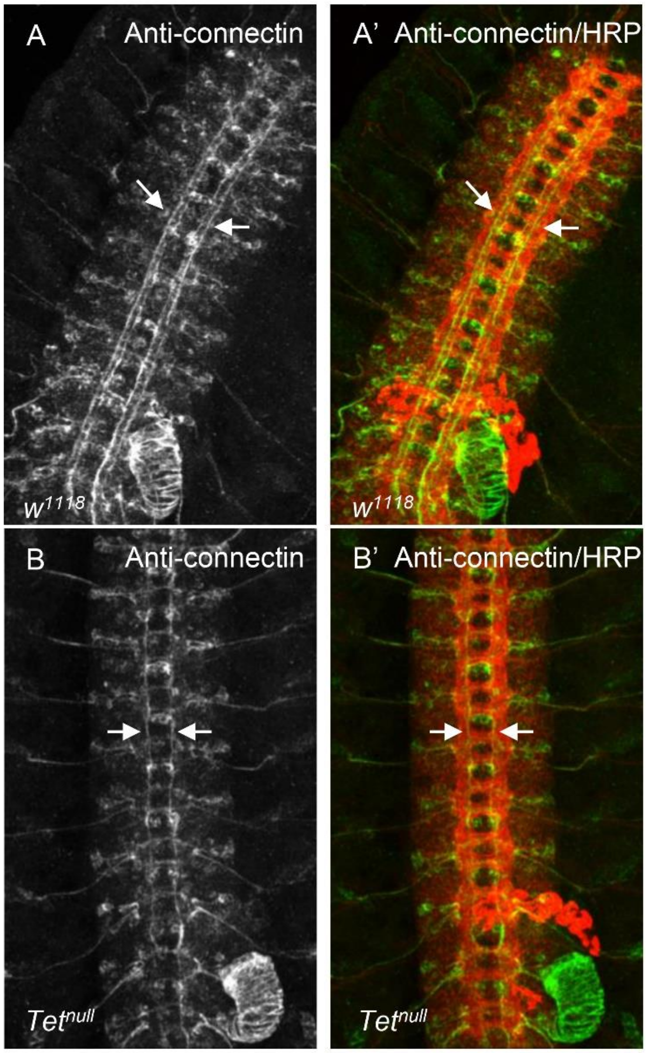


**S8 Fig. Subpopulation of CNS Connectin positive neurons as shown by antibody staining of stage 16/17 embryonic ventral nerve cords**. In wt (**A**) and T*et*$^{null}$/*Tet*$^{null}$ (**B**). **A',B'** show overlays of Connectin+ neurons (green) with the general neuronal marker HRP (red) in *wt* and *Tet*$^{null}$, respectively. Two Con+ tracks run within the longitudinal neuropil on each side of the midline of wt embryos **(A, arrows)** whereas Con+ neurons are present in only one medial track in *Tet*$^{null}$ embryos **(B, arrows).**

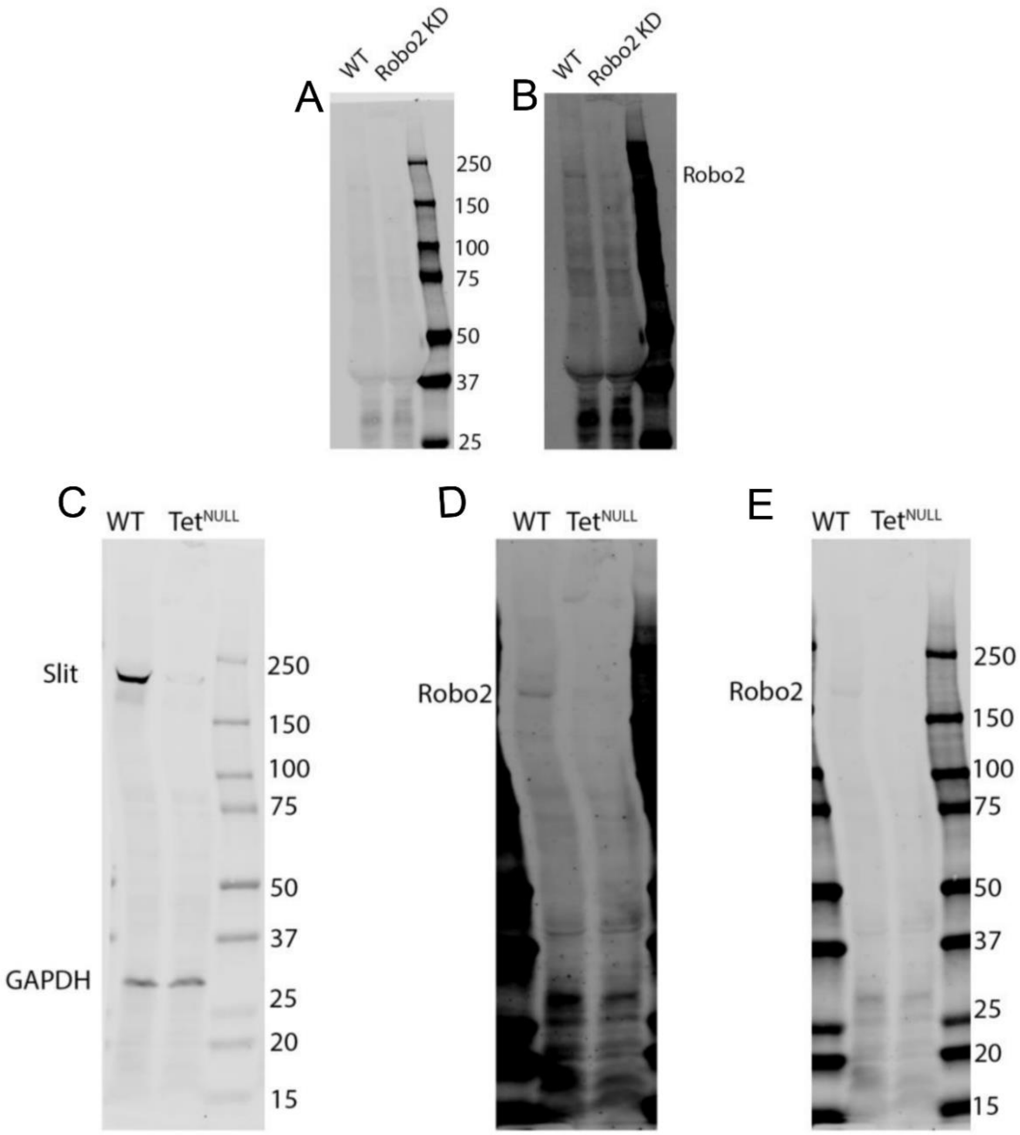


**S9 Fig. Full western blots**: (**A-B).** To test the specificity of the anti-Robo2 antibody we collected 0-12 hour embryos of wt control and *robo2* RNAi knockdown (KD). The wt lane shows a band between 150 and 250 kDa (comparing B to the same western blot in A with low intensity for ladder visualization) which is reduced in the *robo2* KD lane indicating that is robo2 band and specificity of the Robo2 antibody. **C.** wt and $Tet^{null}$ 3rd instar larval brain extracts probed with anti-Slit antibody. **D.** wt and $Tet^{null}$ 3rd instar larval brain extracts probed with anti-Robo2 antibody. **E.** Same western blot in **D.** with low intensity for ladder visualization.

| Genotype | Total # of Hemisegments | FAS2+ Axon Midline Crossing/Hemisegment | FAS2+ Lateral Tract absent/disrupted |
|---|---|---|---|
| $w^{1118}$ | 322 | <1% | 4% |
| $tet^{null}/tet^{null}$ | 298 | 32% | 48% |
| $robo2^{x123}/robo2^{x123}$ | 254 | 22% | 25% |
| $robo2^{x123}/+$ | 259 | 2% | *nd* |
| $robo2^{x123}/+$ ; $tet^{null}/tet^{null}$ | 272 | 30% | *nd* |
| $sli^{2}/+$ | 210 | 1% | *nd* |
| $sli^{2}/+$ ; $tet^{null}/tet^{null}$ | 265 | 48% | *nd* |
| $robo^{z570}/+$ | 111 | 2% | *nd* |
| $robo^{z570}/+$ ; $tet^{null}/tet^{null}$ | 186 | 28% | *nd* |

**S1 Table. Ventral Nerve cord development defects occurring in *tet*$^{null}$, slit, and *robo2* embryos.** Embryos prepared according to Materials and Methods were analyzed for midline crossing of Fas2+ neurons and the presence and integrity of the most lateral longitudinal Fas2+ and Connectin+ tracts.
